# Positive effects of projected climate change on post-disturbance forest regrowth rates in northeastern North American boreal forests

**DOI:** 10.1101/2022.11.29.518357

**Authors:** Victor Danneyrolles, Yan Boucher, Richard Fournier, Osvaldo Valeria

## Abstract

Forest anthropogenic and natural stand-replacing disturbances are increasing worldwide due to global change. Many uncertainties regarding the regeneration and growth of these young forests remain within the context of changing climate. In this study, we investigate the effects of climate, tree species composition, and other landscape-scale environmental variables upon boreal forest regrowth following clearcut logging in eastern Canada. Our main objective was to predict the effects of future climate changes upon post-logging forest height regrowth at a subcontinental scale using high spatial resolution remote sensing data. We modeled forest canopy height (estimated from airborne laser scanning [LiDAR] data over 20-m resolution virtual plots) as a function of time elapsed since the last clearcut along with climatic (i.e., temperature and moisture), tree species composition, and other environmental variables (e.g., topography and soil hydrology). Once trained and validated with ∼240,000 plots, the model that was developed in this study was used to predict potential post-logging canopy height regrowth at 20-m resolution across a 240,000 km2 area following scenarios depicting a range of projected changes in temperature and moisture across the region for 2041-2070. Our results predict an overall beneficial, but limited effect of projected climate changes upon forest regrowth rates in our study area. Stimulatory effects of projected climate change were more pronounced for conifer forests, with growth rates increasing between +5% and +50% over the study area, while mixed and broadleaved forests recorded changes that mostly ranged from -5% to +35%. Predicted increased regrowth rates were mainly associated with increased temperature, while changes in climate moisture had a minor effect. We conclude that such gains in regrowth rates may partially compensate for projected substantial increases in fire activity and other natural disturbances that are expected with climate change in these boreal forests.

## Introduction

Forest natural disturbances are increasing worldwide due to climate change (e.g., fire, drought; Seidl et al. 2017), while land-use-related disturbances such as logging are also intensifying in many parts of the world (Hurtt et al. 2020). Regeneration of severely disturbed forests is undoubtedly among the most powerful drivers of current and future forest dynamics (Danneyrolles et al. 2019; McDowell et al. 2020; Seidl & Turner 2022). These young regenerating forests thus play an increasing role in ameliorating various problems, such as reaching a balance in global carbon dynamics (Pugh et al. 2019; Cook-Patton et al. 2020) and maintaining other forest ecosystem services (Seidl et al. 2016). Yet, to which extent the projected changes in temperature and moisture will positively or negatively affect the ability of these young forests to regenerate and regrowth remains highly uncertain (Anderson-Teixeira et al. 2013; McDowell et al. 2020; Seidl & Turner 2022). These uncertainties are exacerbated in boreal forests, which exhibit more frequent and intense disturbances and slower growth rates than other forest biomes (Gauthier et al. 2015; Seidl et al. 2017; Cook-Patton et al. 2020). Moreover, these northern latitude forests face higher rates of increasing temperature than any other forest on Earth (IPCC 2021). In this context, insights that are based upon new growth data and more reliable models could improve our understanding and predictability of boreal forest regrowth after stand-replacing disturbance.

After a stand-replacing disturbance, several forest ecosystem characteristics and external environmental drivers may control forest regrowth. First, height growth rates are generally maximal in the early stages following a stand-replacing disturbance and tend to decline progressively with stand age as the trees attain their maximum height (e.g., Ryan et al., 2004). Second, tree species composition can strongly modify regrowth due to inter-specific variability in growth rates and maximum tree height. For example, in North American boreal forests, conifer species generally grow more slowly and reach a shorter maximum height than broadleaved species (e.g., Chen & Popadiouk 2002; Messaoud & Chen 2011). Third, regional climate gradients exert considerable control on tree height growth rates. In boreal forests, temperature is the major limitation to growth rates and to maximum height that is reached by the trees (Boisvenue & Running 2006; Huang et al. 2010; Messaoud & Chen 2011; D’Orangeville et al. 2016, 2018). As such, projected increases in temperature for the next few decades should mainly result in net gains in potential boreal forest regrowth (D’Orangeville et al. 2018; Pau et al. 2021; Wang et al. 2022). Fourth, many studies have found that precipitation and climate moisture also significantly influence boreal forest growth, the effects of which can be either positive or negative depending upon interactions between the timing of precipitation, temperature and tree species composition (e.g., Hellmann et al. 2016; D’Orangeville et al. 2016; 2018; Oboite & Comeau 2019). Assumptions regarding the effects of projected changes in precipitation on boreal forest growth thus remain highly uncertain. Last, landscape and site characteristics interact with tree species composition and the regional climate gradient to induce substantial differences in growth rates at the landscape scale. For example, topography results in a high diversity of local temperatures (Nicklen et al. 2016). Topography, surface deposits, and drainage also strongly control site moisture conditions and soil organic content, with mesic slopes generally leading to better growth rates when compared to poorly drained soils with high organic accumulations at lower slope positions (McKenney & Pedlar 2003; Lavoie et al. 2007; Laamrani et al. 2014; Bour et al. 2021).

In this study, we investigated the effects of climate, tree species composition, and other landscape-scale environmental variables upon boreal forest height regrowth following clearcut logging within a large landscape of eastern Canada. Several previous studies have predicted the effects of projected climate changes upon boreal forest growth using tree rings or height data, usually from mature to over-mature trees, without considering their disturbance history or forest age (Huang et al. 2010; D’Orangeville et al. 2016, 2018; Pau et al. 2021). Yet, climate sensitivity of mature to over-mature trees to changing climate may strongly contrast with the sensitivity of more vigorous young regenerating trees (Duchesne et al. 2019). Ignoring the central role of stand-replacing disturbances in boreal forest dynamics may introduce substantial bias in growth models and predictions. To our knowledge, no studies have specifically addressed the effects of projected climate changes upon of boreal forest height regrowth after stand-replacing disturbance over such large region. Moreover, these previous studies focused upon coarse spatial resolutions that could not consider fine-grain variation in growth rates linked to local site conditions. In this study, we developed a model to predict the effects of future climate changes upon post-logging forest regrowth at a subcontinental scale, combined with high spatial resolution. We used a promising robust and recently developed modeling method that was based upon time–since-last clearcut, together with airborne laser scanning data (i.e., LiDAR; Bour et al. 2021). Briefly, forest canopy height that was estimated from LiDAR data over 20-m resolution virtual plots was modeled as a function of time that elapsed since the last clearcut logging and recorded climatic conditions, along with tree species composition and other environmental variables (e.g., topography and soil hydrology). The central assumption is that relationships between current regrowth rates and regional climate gradients provide an excellent means of projecting potential changes in growth to increasing temperature and changes in precipitation. Once trained and validated with ∼240,000 plots, we used this model to test two main hypotheses: 1) projected climate change should have beneficial effects on boreal forest growth. 2) Such effects would differ with forest composition (i.e., coniferous vs. broadleaved forests) and local site conditions (i.e., topography and soil hydrology). For that purpose, we predicted potential post-logging forest height regrowth at 20 m resolution across 237,000 km^2^ following different scenarios that depicted a range of projected changes in temperature and moisture across the study area for the period 2041-2070.

## Materials and methods

### Study area

The study area comprises a 237,000 km^2^ territory of boreal forests in the Province of Quebec, eastern Canada (*sensu* Rowe 1972; Robitaille & Saucier 1998; see Fig. 1). Mean annual temperatures ranges about -2 to 4 °C from north to south and climate moisture varies along an east-west gradient, with a rather continental climate to the west (800-1000 mm of annual total precipitation) and a moister oceanic climate to the east (1000-1200 mm of annual total precipitation; Fig. S1). The east-west gradients include differences in topography and surface deposits, with smooth slopes and abundant lacustrine clay and organic accumulation to the west, and more rugged relief to the east, with a higher proportion of glacial till and rocky deposits (Fig. S2). Black spruce (*Picea mariana*) and balsam fir (*Abies balsamea*) coniferous species tend dominate across the whole boreal zone, with occasional presence of paper birch (*Betula papyrifera*) and trembling aspen (*Populus tremuloides*), together with coniferous white spruce (*Picea glauca*) and jack pine (*Pinus banksiana*; Fig. S3). Further to the south of the study area, within the temperate zone, warm-adapted broadleaved species such as sugar maple (*Acer saccharum*) and yellow birch (*Betula alleghaniensis*) dominate the landscapes, which contain scattered patches of boreal forest (i.e., dominated by either black or white spruce, balsam fir, jack pine, trembling aspen or paper birch).

**Fig. 1.**
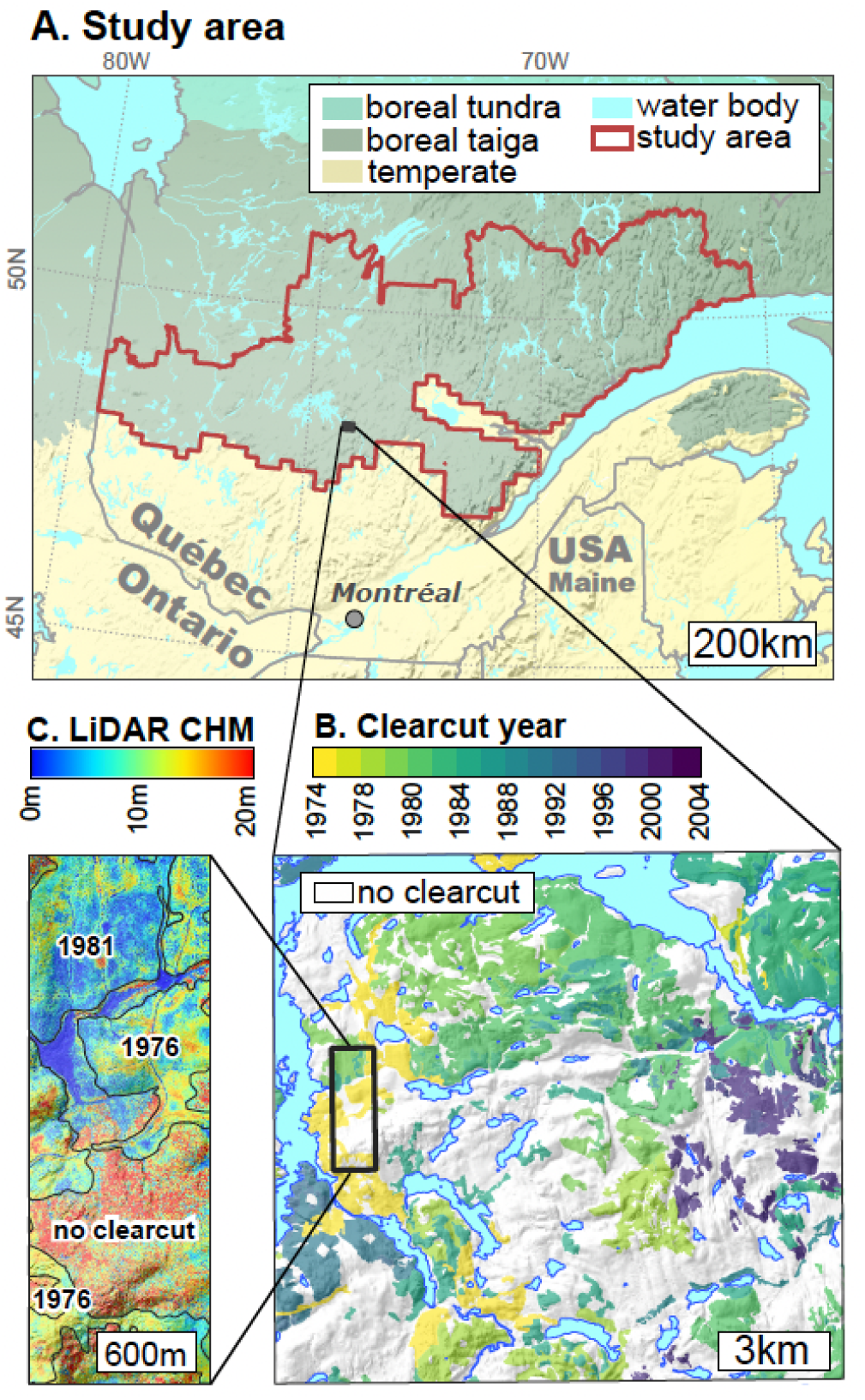
Study area (A) and illustration of time-since-last clearcut data data (B) and forest canopy height data (i.e., LiDAR canopy height model [CHM]; C). Virtual 20 × 20 m plots were selected in clearcut areas across the whole boreal zone (delimited by the red line) and only within stand that were dominated by boreal species in the temperate zone of southern Quebec. Prediction of climate-induced changes in forest regrowth were limited to the study area (delimited by the red line).

Stand replacing disturbances, mostly fire and logging, are widespread within the study area (Boucher et al. 2017). According to the Quebec government’s forestry maps (MFFP 2018), between 1980 and 2015, ∼21% of the forest cover of the study area was clearcut logged, and ∼5% burned (i.e., 0.58% and 0.15% per year, respectively). About a quarter of the forests logged since 1980 were replanted, while the rest regenerated naturally. In this study, we concentrated our analyses on clearcut disturbance rather than fire since the annual area affected by logging is more homogeneously distributed in space and time and then much more adapted for developing a space-for-time modeling approach.

### Time-since-last clearcut logging data

We extracted time-since-last clearcut data from the Quebec government’s latest available forestry maps (MFFP 2018). The polygons that were drawn at the 1:20 000 scale (with a minimum 4 ha area) are based upon the interpretation of high-resolution aerial photographs and annual harvesting reports (Fig. 1). Each polygon informs on the type and year of the last known stand-replacing disturbance since the 1950s (Fig. 1), along with the dominant species of the stands. We randomly sampled hundreds of thousands of virtual LiDAR 20 m × 20 m plots within disturbed areas of our study area. We also sampled plots in disturbed areas in the temperate zone southern to the study area (Fig. 1), but only within stands of forests that were dominated by boreal species (i.e., either black or white spruce, balsam fir, jack pine, trembling aspen, or paper birch). We retained these southern parcels for incorporation into our model because these boreal forests are currently growing under warmer climate conditions, which likely correspond to the projected climate conditions in the boreal zones of the future (Fig. S1). To the north, the delimitations of the study area were defined by the availability of LiDAR data (most forests to the north of our study area are not exploited and thus not surveyed in Quebec’s government provincial forest inventory program).

According to the forestry maps, all plots that were retained belonged to two clearcut logging categories: clearcut with natural regeneration; and clearcut with tree planting. The plots that were selected were at least 50 m from the clearcut polygon boundaries to avoid edge effects and stand margin delineation errors (see Bour et al. 2021, for details). The age of each 20 m × 20 m plots within the clearcuts (i.e., time-elapsed-since last disturbance) was then calculated as the difference between LiDAR acquisition year (see details below) and logging or planting year. We only retained plots that were aged > 10-years-old, given that trees < 10-years-old can be confused with ericaceous shrubs or alder (*Alnus* spp.) (Matasci et al. 2018). The oldest recorded plots were 70-years-old, given that no reliable logging history data were available prior to 1950. Precommercial thinning is a widespread silvicultural treatment in the boreal forests that reduces stand density and competing vegetation (Ashton & Kelty 2017), which is occasionally applied a few years following post-logging natural regeneration or planting in our study area. In our dataset, 15% of naturally regenerated stands and 3% of plantations were treated using precommercial thinning. Consequently, we considered four distinct types of silvicultural treatment for the following growth analyses: clearcutting with natural regeneration alone (72% of the 238,519 plots in the final dataset); clearcutting with natural regeneration + precommercial thinning (15%); clearcutting with planting (10%); and clearcutting with planting + precommercial thinning (3%).

### Canopy height data

Forest canopy height was determined using airborne LiDAR data that the Quebec government acquired as part of its provincial forest inventory program. For the data that were used in this study, aerial surveys were realized between 2012 and 2019, with point densities ranging from 1.5 to 8 pulses.m^-2^ (median = 3 pulses.m^-2^). Raw point clouds were transformed by analysts from the Quebec government into digital elevation and canopy height models (DEM and CHM, respectively, both at 1-meter resolution; Fig. 1C). To do so, raw point clouds were first classified into ground and non-ground returns and a DEM was fitted to the ground returns to produce a 1-m resolution raster. The DEM was subtracted from elevations of all non-ground returns to produce a normalized point cloud. Finally, the height of each 1 m pixel of CHM was defined as the highest return of the normalized point cloud.

The 1-meter resolution CHMs were aggregated into the 20 m × 20 m plots simply by assimilating the height of each plot as the 95^th^ percentile (P95) of its CHM pixel heights and after removing pixels < 1 m in height. The 95^th^ percentile is frequently used to produce CHM from raw point clouds (White et al. 2013), and exclusion of the lowest returns (< 1 m) is usually applied to remove returns attributable to the herbaceous-shrubby ground vegetation (Nyström et al. 2012). We also compared this LiDAR-based metric (P95) with field-based measurements of forest canopy height using 249 forest inventory plots that were surveyed in the field during the same year as the LiDAR survey (Fig. S4). This comparison showed that P95 is very closely related to the mean height of dominant and co-dominant trees that are measured on the field (Pearson *r* = 0.89). To avoid spatial autocorrelation in this forest height measure, we only retained sample plots in clearcuts or planted areas that were separated by a minimum distance of 250 m, a threshold that was validated with a semi-variogram (Fig. S5).

The 1:20 000 polygons that identified disturbed areas have a minimum area of 4 ha. These polygons can include small patches of remnant forest that have not been harvested (e.g., plots of 20 m × 20 m = 0.04 ha). If such remnant forest patches exist, their occurrence can incur bias in the analysis. To remove this potential bias, we excluded plots in which we identified aberrant heights for a given age from the LiDAR data. The threshold values for identifying aberrant heights were defined using a database of nearly a million trees, from which age and height have been measured within our study area through the provincial forest inventory programs since 1981 (i.e., individual tree field-based height measurements and tree-ring age estimates). Threshold values were estimated separately for the six main species of our study area. To do so, we first calculated the maximum height threshold for a given age as defined by the 99^th^ percentile of field-measured tree height (Fig. S6). The maximum height threshold per age class and per bioclimatic domain (i.e., part of the Quebec’s government ecological classification system; Robitaille & Saucier 1998) was then defined as the mean of individual species threshold weighted by their relative abundance in each bioclimatic domain (Fig. S6). This final selection excluded ∼10,000 plots (∼4% of the dataset before filtering), and the final dataset was comprised of 238,519 plots.

### Topographical, climate and forest composition data

We extracted seven explanatory variables that could influence canopy height growth (Table 1). First, we extracted the proportion of broadleaved trees at the center of the 20 m × 20 m plots from a 30-meter raster product that was made available by the Canadian Forest Service (Fig. S7) The proportion of broadleaved trees has been modeled with Landsat images and is based upon information from more than 10,000 inventory plots (for a more detailed explanation of the methodology, see Guindon et al. 2020). Second, we extracted several variables from climate gridded datasets (McKenney et al. 2011) at a 300-arcsecond resolution (∼55 km^2^) that contain monthly climate observations since 1950 (Fig. S1). For each individual 20 m × 20 m plot, climate normals were calculated as the average of observed variables starting from the year of clearcutting and stopping at the year of LiDAR acquisition. We calculated the yearly mean daily temperature (T_MEAN_, °C) and maximum daily temperature (T_MAX_, °C). We calculated the mean annual Climate Moisture Index (CMI), which is defined, for a given period, by the sum of total precipitation minus the sum of potential evapotranspiration (the latter was calculated following Baier & Robertson, 1965). CMI was calculated for the growing season (May to September; CMI_GS_) and for the summer period (June to August; CMI_SM_). Finally, we extracted three variables from LiDAR DEM (1-m resolution) that characterized the topography and soil hydrology of the 20 m × 20 m plots: mean elevation, slope (%), and topographic wetness index (TWI; Sørensen et al. 2006). One other variable that was taken from forestry maps characterized five geological surface deposits (Fig. S2): glacial tills (74% of the entire dataset), lacustrine clays (13%), fluvio-glacial (8%), rocky outcrops (3%) and organic soils (2%).

**Table 1.** Description of variable sources, type (Cont.: continuous), and range in the training and validation datasets. The first number is the mean value in the range column of continuous variables, while numbers within parentheses are minimum and maximum. Climate, topographical and percent of broadleaved trees maps are shown in Fig. S1, S2 and S7 respectively.

| Variables | Source | Type (unit) | Range |
| --- | --- | --- | --- |
| Canopy height (P95) | LiDAR | Cont. (m) | 9.36 (1.46, 22.320) |
| Time since last clearcut | Forestry maps | Cont. (year) | 30 (10, 70) |
| Silvicultural treatment | Forestry maps | Categorical | Clearcut, Clearcut + thinning, Planting, Planting + thinning |
| Percentage of broadleaved trees | Landsat + field plot model | Cont. (%) | 40 (1, 99) |
| Mean annual temperature ($T_{\text{MEAN}}$) | Meteorological | Cont. (°C) | 1.5 (0.54, 6.15) |
| Summer climate moisture ($\text{CMI}_{\text{SM}}$) | Meteorological | Cont. (mm) | -117 (-250, 31) |
| Elevation | LiDAR | Cont. (m a.s.l.) | 432 (3, 1074) |
| Slope | LiDAR | Cont. (%) | 12.6 (0.4 – 125.7) |
| TWI | LiDAR | Cont. (no unit) | 5.61 (0.3 – 23.9) |
| Surface deposit | Forestry maps | Categorical | Glacial tills, Fluvioglacial tills, Glaciolacustrine clays, Rocky, Organic |

### Canopy height growth modeling

We used random forest regression (Breiman 2001) to model canopy height growth since such machine learning approaches are non-parametric and very efficient in modeling non-linear ecological data with complex interactions (Christin et al. 2019). We used the *ranger* function that is included in the *ranger* package (Wright & Ziegler 2017) to train a random forest regression model with 500 trees and three variables to guide the split at each node. We trained the model with a random sampling of 80% of the original dataset (i.e., 190,815 observations), while we held back the remaining 20% (i.e., 47,704 observations) for measuring model predictive performance. The training and validation datasets exhibited identical distributions for all continuous and explanatory variables that were integrated into the model. In the model, canopy height depends upon the following nine explanatory variables: time-since-last clearcut; silvicultural treatment; temperature; climate moisture (CMI); proportion of broadleaved trees; elevation; slope; TWI; and surface deposits. To retain only two climatic variables (temperature and moisture), we compared the model output combining temperature (T_MEAN_, T_MAX_) and climate moisture (CMI_GS_, CMI_SM_) variables. Output with different combinations of temperature and moisture were almost identical (Fig. S8). Consequently, we retained yearly mean daily temperature (T_MEAN_) and CMI from June to August (CMI_SM_) because these two variables exhibited the lowest correlation (Pearson *r* = -0.33), which could help to separate the effects of temperature and moisture on canopy height growth. Overall, all continuous explanatory variables that were included in the model showed low to moderate correlations with one another (Fig. S9).

### Canopy height growth rate projections across the boreal zone

Once trained, the model was used to produce maps of predicted post-logging forest canopy height growth at 20-meter resolution across the boreal zone (Fig. 1) according to 27 scenarios. The 27 scenarios represent all possible combinations of three conditions for T_MEAN_ (baseline, representative concentration pathways RCP 4.5, and RCP 8.5), three for CMI_SM_ (baseline, RCP 4.5, and RCP 8.5), and three for cover composition types (coniferous, mixed, broadleaved).

T_MEAN_ and CMI_SM_ baseline values were defined from the 40-year normals that were recorded during the 1980-2020 period. Projected T_MEAN_ and CMI_SM_ for the RCP 4.5 and 8.5 emission scenarios correspond to the 2041-2070 normals that were obtained as median values across 11 global circulation models (Fig. S1; all climate simulations with different models and scenarios were taken from the non-profit OURANOS organization on climate change science, see https://www.ouranos.ca). In brief, RCP 8.5 represents a high-end climate scenario with continuous high greenhouse gas emissions, while RCP 4.5 is a middle-of-the-road scenario (IPCC 2021). These correspond to an increase in T_MEAN_ of about 2°C and 3°C for the RCP 4.5 and 8.5 scenarios, respectively, where increases are relatively uniform across the study area. Projected changes in CMI_SM_ were more spatially contrasted, with a major portion of the study area experiencing more humid climate, while other portions (towards the west) experienced slightly drier climate under both RCP 4.5 and 8.5 scenarios (Fig. S1). Almost all of these projected future temperatures and climate moisture conditions lie within the range of climatic conditions that are used to train the random forest model (i.e., calibration range; see Fig. S1). To understand how climate change may modify the growth rates of different forest cover composition types, we also set three distinct possibilities with forest cover compositions being fixed to coniferous, mixed, or broadleaved types across the study area (i.e., 5%, 50% and 95% broadleaved trees, respectively). In all scenarios, permanent topographical variables (elevation, slope, surface deposits and TWI) remained unchanged across the study area. Since silvicultural treatments had a marginal effect on modeled height growth (Fig. S10), we assigned all scenarios with the most common silvicultural treatment (i.e., clearcut alone; 72% of the dataset). For each 20 m × 20 m pixel of the study area and according to these 27 scenarios, we computed potential growth as the predicted height at 50 years, averaged at an annual rate to obtain a map of canopy growth in cm.year^-1^. We chose potential height at 50 years, given that this metric is widely used in forestry (commonly referred as “site index,” e.g., Messaoud & Chen 2011; Pau et al. 2021), given that most boreal forests of our study region reach a height growth plateau at this age (see *Results* section). We also computed predicted uncertainty using the quantile random forest regression approach (Meinshausen & Ridgeway 2006) in which the variance of predicted values is quantified between trees within the random forest model and used as a metric of prediction uncertainty. Note that this corresponds to the uncertainty linked to the random forest modelling approach but not to the uncertainty related to climate projections.

## Results

The comparison between predicted and observed forest canopy height in more than 47,000 observations from our validation dataset highlighted the strong predictive power of the model (Fig. 2). The model explained 75% of variation in the validation dataset height (i.e., LiDAR P95) with a linear relationship very close to 1:1, together with a relative RMSE = 21% and a mean error = 0.01 m. The importance of the explanatory variables was measured by permutation tests that were performed on random forest out-of-bag samples (i.e., % increase in MSE; Fig. 2). Our results show that time-since-last clearcut has the most decisive factor influencing canopy height. Under all environmental conditions, canopy heights tended to assume a logarithmic function of time-since-last clearcut, with the highest growth rates being observed during the first few decades, after which a plateau was reached at about 50 years following clearcutting (Fig. 3 and S10). The second most influential variable was T_MEAN_ (temperature). Although the weightings of these first two variables dominate, in decreasing order of importance, the other prominent variables were the percentage of broadleaved trees and CMI_SM_ (Fig. 2). These four variables showed notable interaction effects with one another. Temperature had a significant positive effect on growth rates under all moisture conditions and forest cover composition types (Fig. 3). Conifer-dominated forests tended to display lower growth rates than mixed and broadleaved forests under all temperature and moisture conditions. These differences were particularly marked in cold and moist sites and disappeared in more hot and dry sites (Fig. 3). Climate moisture (CMI_SM_) negatively affected forest growth, particularly in conifer stands. In mixed and broadleaved forests, such effects were attenuated (Fig. 3). Topographical variables had a secondary influence on canopy height (Fig. 2). Globally, the best growth rates were found in lower elevation and moderate slopes that were associated with low-moisture glacial, fluvio-glacial or clay deposits (Fig. S10). In comparison, lower growth rates were found in high altitude sites, high moisture flat sites that were associated with organic deposits, or on rocky deposits (Fig. S10). Finally, silvicultural treatments had a very limited effect on forest growth compared to other variables that were included in the model (e.g., climate, topography; Fig. 2; Fig. S10).

**Fig. 2.**
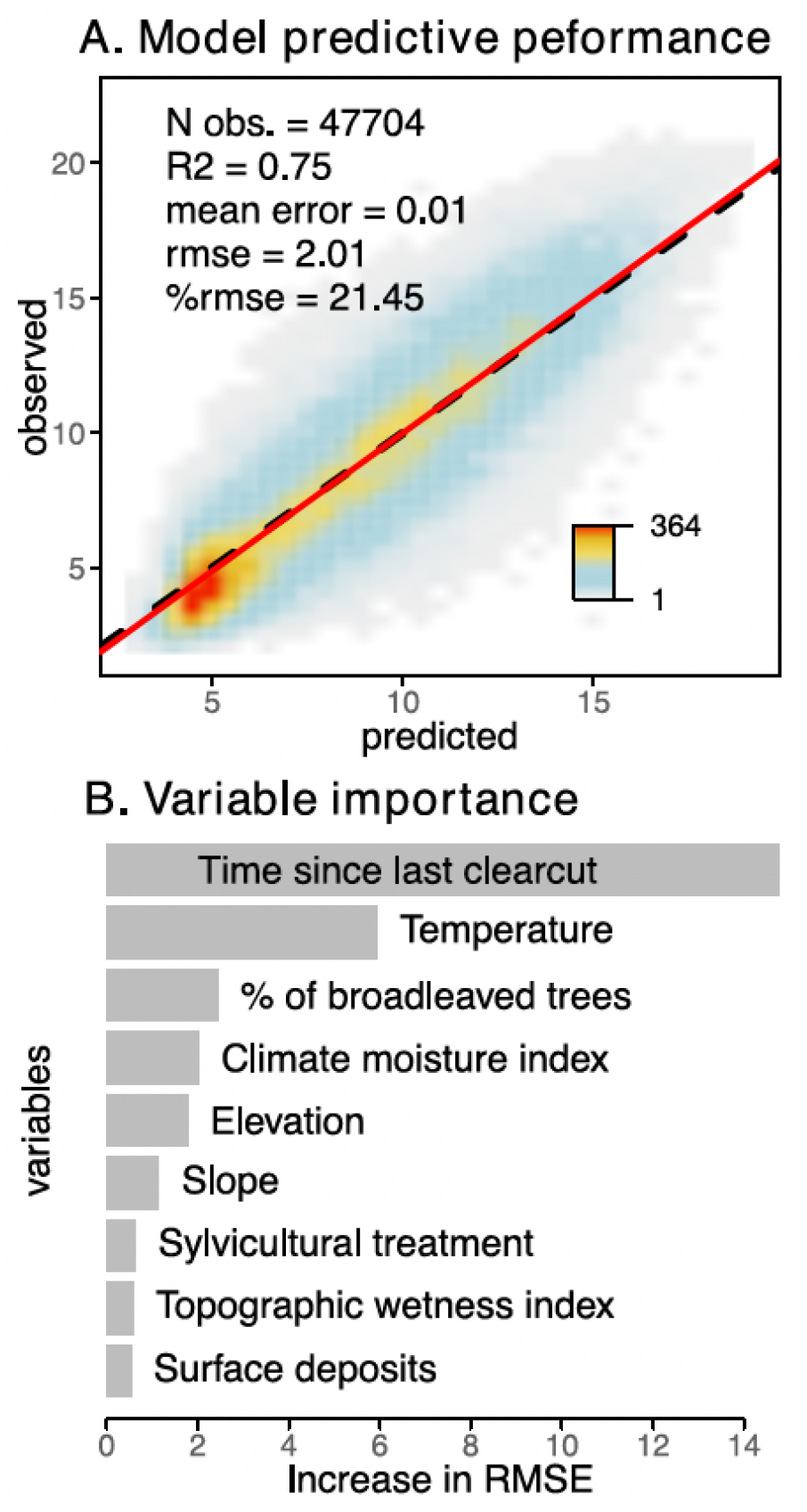
Model predictive performance (A) and variable importance (B). Model predictive performance in (A) was evaluated by comparing observed and predicted forest heights (P95) in the validation dataset (20% of the entire dataset; *N* = 47 704). Point cloud density in (A) is displayed as a color gradient. Variable importance in (B) was evaluated through the impact of single variable permutation upon model predictive performance (i.e., increase in RMSE).

**Fig. 3.**
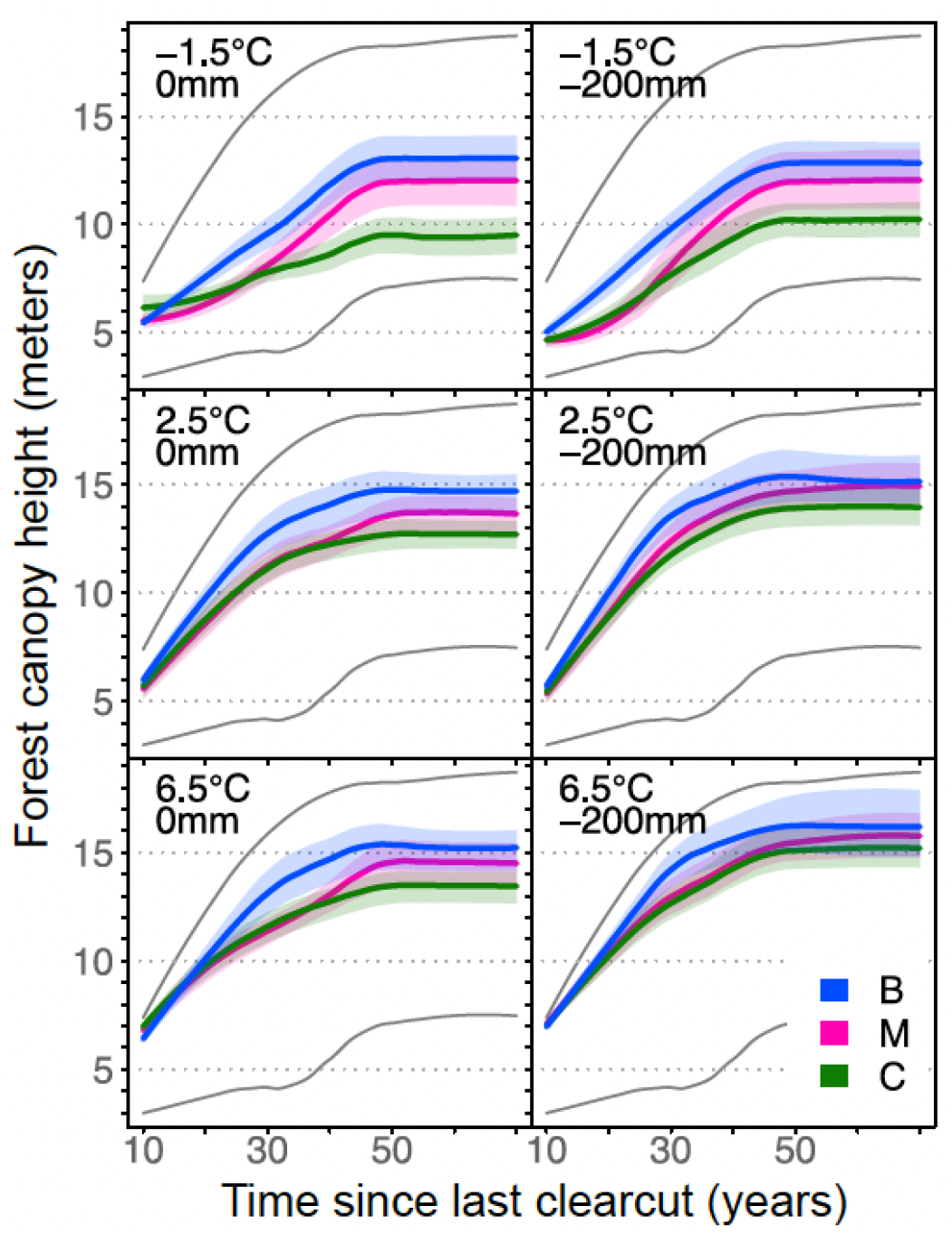
Interactive effects of time-since-last clearcut, forest composition, temperature, and climate moisture variables upon forest height. A unique random sampling of 999 observations in the original dataset was used to predict forest canopy height across all one-year time-since-last clearcut classes. The fine grey lines show the whole variability of predicted heights without fixing any parameters. Colored lines and ribbons show the central tendency and 70% confidenc intervals of predicted heights after fixing different parameters. Forest composition was fixed a coniferous, mixed, or broadleaved (respectively: C = 5%, M = 50%, and B = 95% of broadleaved trees). Climate variables (T_MEAN_ and CMI_SM_) were fixed within their observed range in the original dataset and fixed values are shown in the upper left corners.

During the first 50 years post-logging, projected potential canopy height regrowth rates across the study area following the 27 baseline scenarios (i.e., 1980-2020 climate) ranged from 15 to 35 cm.year^-1^ (i.e., heights between 7.5 and 17.5 meters at 50 years, respectively; Fig. 4). Conifer-dominated forests showed growth rates ranging from 15 to 25 cm.year^-1^ in > 75% of the study area (i.e., between 7.5 and 12.5 meters at 50 years, respectively; Fig. 4), while median growth rates of mixed and broadleaved forest were ∼27 cm.year^-1^ (i.e., ∼13.5 meters at 50 years; Fig. 4). Simulated changes in regrowth rates with different climate change scenarios (i.e., projected climate for the 2041-2070 period) mainly benefited all forest cover composition types (Fig. 4). Increased regrowth rates were mainly associated with increased temperature, while projected changes in climate moisture had a minimal or even insignificant effect on forest regrowth (Fig. S11). The stimulatory effects of projected climate change were more pronounced for conifer forests, with growth rates increasing between 5% and 50% over almost the entire study area in the RCP 8.5 scenario compared to the baseline (Fig. 4). In comparison, most mixed and broadleaved forests recorded net changes ranging between -5% and 35% under the RCP 8.5 scenario (Fig. 4 and S12). This larger effect of increased temperature upon conifer forests reduced the initial differences in growth rates between conifers and mixed or broadleaved forests (Fig. S13). The sites with lowest baseline growth rates (i.e., 15 to 20 cm.year^-1^) also recorded the largest proportional increases in growth rates with increased temperatures (Fig. 4).

**Fig. 4.**
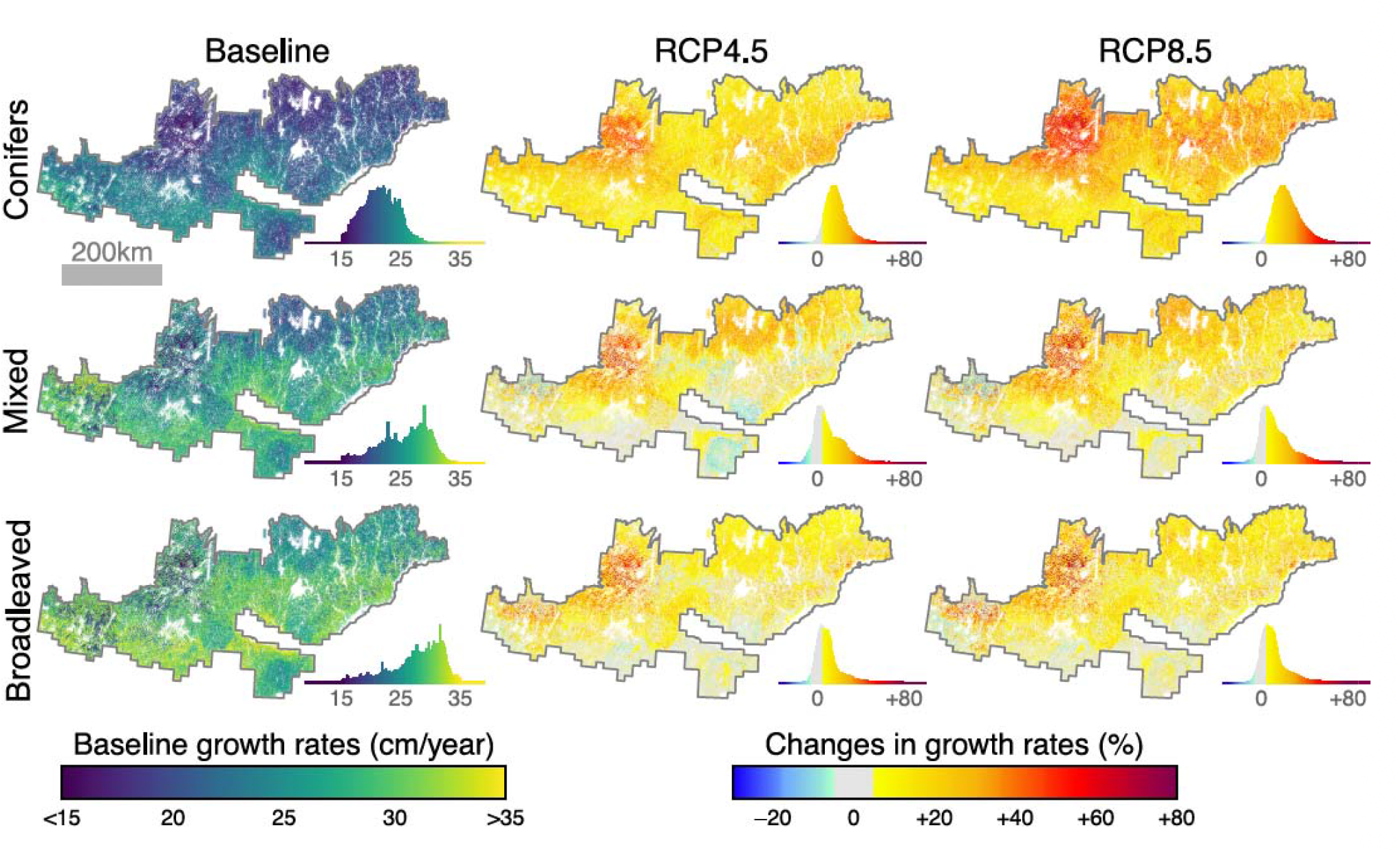
Predicted potential post-logging regrowth rates within the boreal zone. For each forest composition classes (i.e., A: coniferous, B: mixed, or C: broadleaved), height growth rates in the first 50 years following clearcut was predicted following different climate scenarios (see Fig. S1). Temperature (T_MEAN_) and moisture (CMI_SM_) baselines (1980-2020) are shown in the first column and displayed as growth rates. Changes in height growth rates with projected temperature and moisture for the RCP4.5 and RCP8.5 scenarios (2041-2070) are displayed a percentage of changes from baseline scenarios. All combinations of effects individual changes in temperature and moisture for baseline, RCP4.5, and RCP8.5 scenarios are shown in Fig.s S11, S12 and S13.

## Discussion

Our modeling approach that was based on airborne LiDAR and time-since-last clearcut data had great power to predict forest regrowth rates after logging in a large landscape of northeastern North American boreal forest. This research represents a significant milestone toward a better understanding of climate change effects on boreal forest growth. First, it predicts the effects of climate changes upon boreal forest growth, specifically after stand-replacing disturbance. Previous studies were limited to predicting forest growth without considering disturbance history or forest age (e.g., Huang et al. 2010; D’Orangeville et al. 2016; 2018; Pau et al. 2021). Ignoring such central aspects of boreal forest dynamics may introduce significant bias in model predictions (Duchesne et al. 2019). Second, compared to models alimented with field-based data, our model is based upon remotely sensed data that are less monetarily costly and time-consuming to acquire, while it can accurately predict forest regrowth rates at the sub-continental scale with much finer spatial grain. In any event, the predicted height growth rates of our model are very consistent with those found in previous studies of the boreal forests (i.e., mostly comprised between 10 and 40 cm.year^-1^; Pedlar & McKenney 2017; Oboite & Comeau 2019; Pau et al. 2021).

The regional gradient of temperature normals had a major positive effect on forest height regrowth rates, with height at 50 years varying from ∼10 to 12 m in the coldest conditions to ∼14-18 m in warmer conditions. This is not surprising given well-known major temperature constraints on forest growth at these northern latitudes (Boisvenue & Running 2006). The main underlying mechanism behind the positive temperature-growth relationship is that increased mean annual temperatures imply longer growing seasons (e.g., Huang et al. 2010; D’Orangeville et al. 2016; 2018), thereby allowing trees to grow faster and taller (e.g., Messaoud & Chen 2011; Pau et al. 2021). An additional ecological explanation is that southern and warmer boreal forest stands generally grow more densely than those in colder conditions (Beaudoin et al. 2014). Higher stand density implies higher competition for light, a context in which trees tend to allocate more resources to their growth in height at the expense of radial growth (e.g., McCarthy & Enquist 2007; Schneider et al. 2018). In our methodological context, higher stand density increases the probability of LiDAR detecting high points of the canopy in the 20 m × 20 m plots, which may represent a slight bias in our data.

Compared with temperature, the regional gradient of climate moisture normals had a less marked, but nevertheless negative effect on forest regrowth. The effect of climate moisture also substantially interacted with forest composition, since it was more significant for conifer-dominated forests. Such negative effects of climate moisture on boreal tree species growth have already been reported in North American and Eurasian boreal or cold mountain forests (D’Orangeville et al. 2016; 2018, Hellmann et al. 2016; Buechling et al. 2017; Pau et al. 2021). This counterintuitive negative effect of climate moisture may be linked to several non-exclusive explanations. First, our study area represents one the wettest boreal regions in the world (annual total precipitation ranging from 800 to 1200 mm versus 250 to 700 mm for most western North American or Eurasian boreal forests; D’Orangeville et al. 2016). In these rather wet forests, the high snowpack meltwater may ensure sufficient soil water levels for most of the growing season, thereby making growing season climate moisture a non-limiting factor for forest growth (D’Orangeville et al. 2018). Second, in such context, high growing season precipitation could also incur indirect reduction in forest growth through a decrease in solar radiation and soil temperature (Bergeron et al. 2007). Yet, previous studies have found that the effect of climate moisture on forest growth became positive after crossing a threshold of warm temperatures, which contrasts with our results (D’Orangeville et al. 2018; Pau et al. 2021). This divergence could be explained because our model specifically examines young regenerating stands, which may require less moisture to support their growth than would older stands with larger living biomass and high related water demand (Duchesne et al. 2019). For example, young northern temperate pine (*Pinus strobus*) plantations have been found to use water more efficiently than older ones that exhibited higher evapotranspiration levels during drought events (Skubel et al. 2015).

Two principal limits to our approach could also constrain our ability to properly model the effects of climate variables upon forest height growth. First, our model accurately predicted well-known differences among forest composition cover types with slower growth rates for coniferous (i.e., black spruce, balsam fir, and jack pine) compared to broadleaved forests composed of fast-growing, shade-intolerant trembling aspen and paper birch (e.g., Chen & Popadiouk 2002). However, our data only measure forest composition at a functional level (i.e., conifer-broadleaved ratio), but cannot separate forest compositions at the species level. Obviously, the climate gradient in our study area is linked to tree species composition (see Fig. S3): black spruce dominates the coldest northern sites, while balsam fir and white spruce can represent a major proportion of stands in warmer southern sites. The driest portions of our study area also show the greatest proportions of trembling aspen and jack pine. Since different species within the conifer or broadleaved functional groups broadly diverge in terms of growth rates and climate-growth relationships (D’Orangeville et al. 2018; Marchand et al. 2019), some variation in height growth due to species composition is implicitly accounted for in the modeled effects of climate gradients. On an even finer phylogenetic scale, our model does not account for local genetic adaptation within the same species, which may significantly influence tree sensitivity to climate variables (e.g., Depardieu et al. 2020; Girardin et al. 2021). The second limit of our approach lies in the use of climate normals that cannot model the effect of single-year climate extremes on forest growth. However, monthly or yearly events such as extreme summer heat, drought events, or spring frost can have significant and long-lasting effects on forest growth (Marquis et al. 2020).

Regardless of these limits, our simulations with projected changes in climate normals for the 2041-2070 period resulted in an overall increase in potential post-logging forest regrowth. These results are consistent with previous research in our study area (i.e., mostly between -5% and +50%; D’Orangeville et al. 2018; Pau et al. 2021), although another recent study found higher near-term increases (i.e., +40 to +52% on average across Canada’s boreal ecozone for the 2050s time period; Wang et al. 2022). In our results, beneficial effects are pervasive across the study area, but are more pronounced for conifer forests and for cold northern, slow-growing forests, which agrees with these studies to some extent. Our simulations show that increases in potential post-logging growth rates are strictly due to increased temperatures for the most part, because projected changes in climate moisture have virtually no effect (Fig. S11). This response contrasts with a previous study, which showed that the beneficial effects of increased temperature on boreal forest growth could reverse after crossing a threshold of warmer temperature (about +3 to 4 °C), beyond which a negative impact may occur with an associated decrease in climate moisture (D’Orangeville et al. 2018). This is likely linked to the specific nature of our study that models the growth rates of young, vigorous regenerating trees, rather than the growth of mature to over-mature trees that may be much more sensitive to climate moisture effects (Duchesne et al. 2019).

## Conclusions

Our results show an overall beneficial effect of the projected increase in temperature on forest height regrowth in the 50 years following a disturbance in northeastern North American boreal forests. These results provide new knowledge and insights into the effects of future climate change on these forests, which are necessary for sustainable forest management and global change assessments. A marked increase in fire activity and other natural disturbances is expected due to climate change in these boreal forests (e.g., Boucher et al. 2018; Coogan et al. 2019; Seidl et al.,2020). Previous studies modeling the effects of climate change upon forest growth have argued whether their projected changes would be either insufficient (D’Orangeville et al. 2018; Pau et al. 2021) or sufficient (Wang et al. 2022) to compensate for effects of future natural disturbances. Yet, these previous studies modeled forest growth without considering disturbance history or forest age, and in some cases, even used mature to over-mature trees as reference data. By predicting the effects of climate change upon boreal forest growth specifically after stand-replacing disturbance, our results may be more appropriate to pronounce on these issues. Gauthier et al. (2015) have determined that it would take an increase in growth rates of at least 50% on average to offset the loss of biomass due to projected future burned areas alone. Despite growth gains being a pervasive outcome of our results, they remain limited (i.e., net changes range mostly between -5% to +50%), suggesting that growth gains may partially compensate for the projected effects of disturbances, but this should not allow any increase in harvested volumes. How climate change combines with disturbance regimes to affect boreal forest growth, functioning and management remains a central topic for future research.

## Acknowledgements

We thank Batistin Bour, Luc Guindon, Jason Laflamme, Jean-François Bourdon, Martin Riopel, Steve Bedard and Michel Campagna for their support and advice concerning forest-related data. We also thank Pascal Bourgault and Travis Logan for their help in accessing climate data and William F.J. Parsons for carefully editing the manuscript. Funding was provided by the *Contrat de service de recherche forestière* number 42332166-1 obtained by R. Fournier of the Université de Sherbrooke from the Ministère des Forêts, de la Faune et des Parcs (Quebec, Canada). This project also benefited from research grant number 566938 - 21 from the Alliance program of the Natural Sciences and Engineering Research Council of Canada (NSERC).

## Data Accessibility Statement

All raw forest data used in this study are freely available at https://www.donneesquebec.ca/ (i.e., LiDAR canopy height models, time-since last clearcut data and other environmental variables). Raw climate data are freely available at https://www.ouranos.ca. The assembled virtual plots dataset and R codes used to train the random forest model are available at https://doi.org/10.6084/m9.figshare.20418792.v1. Conditional on manuscript acceptance, the results produced in this study (i.e., predicted potential post-logging regrowth rates maps shown in Figure 4) will be freely available online.

