## Supplementary Information for "Positive effects of projected climate change on post-disturbance forest regrowth rates in northeastern North American boreal forests"

**Figure S1.** Baseline and projected changes in yearly mean daily temperature ($T_{MEAN}$ in °C) and summer climate moisture index ($CMI_{SM}$ in mm) within the study area. Baselines correspond to observations for the 1980-2020 period, while projected changes with the RCP4.5 and RCP8.5 scenarios correspond to the 2041-2070 period. $CMI_{SM}$ is defined by the sum of summer (June to August) total precipitation minus the sum of potential evapotranspiration (the latter is calculated following Baier & Robertson, 1965). For both $T_{MEAN}$ and $CMI_{SM}$, difference with baseline is shown below RCP4.5 and RCP8.5 scenarios. Distributions of $T_{MEAN}$ and $CMI_{SM}$ observed in the training dataset and in Baseline, RCP4.5 and RCP8.5 scenarios for the boreal zone (delimited in by a red line on the maps) are shown beneath baseline maps.

**Figure S2.** Maps of elevation (A) and surface deposits (B) within the study area. Elevation was derived from LiDAR data (1-m resolution; https://www.donneesquebec.ca) and surface deposit categories were taken from forestry maps (1:20 000 polygons; https://www.donneesquebec.ca; MFFP, 2018). LiDAR-based elevation data at 1-m resolution were used to derive slope and topographic wetness index (TWI; Sørensen et al., 2006) maps across the study area. All topographical variables were finally aggregated at 20 m resolution to project potential post-logging forest regrowth across the boreal zone (delimited by black line).

**Figure S3.** Distribution and height-age relationships of the six main boreal tree species in the field-based Quebec's government forest inventory programs since 1981. Maps were produced as the proportion of total basal area represented by each species at the ecological district-level that are part of the Quebec's government ecological classification. Tree height-age relationships were defined using a database of almost a million trees, the age and height of which have been measured since 1981 (i.e., single tree field-base height measurements and tree-ring age estimates). For each tree species, lines show median and maximum (99$^{th}$ percentile) for a given age (thick and fine lines, respectively). Red lines show the height found in trees within the boreal zone (delineated by red lines on the maps), while black lines show the height found in trees within the temperate zone.

**Figure S4.** Comparison of airborne LiDAR and field-based measures of forest height. The comparison was established using 249 forest inventory plots (400 $m^2$) that surveyed in the field durng the same year as the LiDAR survey (see map A; bioclimatic domains are part of the Quebec's government ecological classification).). We then compared different LiDAR-based metrics with different field-based metrics (B). LiDAR-based metrics were the mean of CHM 1 $m^2$ pixels >1 m in height (LiDAR mean), and the 50$^{th}$, 75$^{th}$, 95$^{th}$ and 99$^{th}$ percentiles of CHM 1 $m^2$ pixels >1 m in height (LiDAR P50, P75, P95 and P99, respectively). Field-based measurements were the maximum height of measure trees (Max height), the mean height of measure trees that were classified as dominant or codominant (Mean dom. height) and the mean height of all measured trees (Mean height). Plot aboveground biomass (AGB) were also estimated with species-specific allometric equations (Lambert et al., 2005). Best correlations were found between LiDAR P95 and the mean of all measured tree heights (see C, red line shows the linear regression between the two measurements). P95 is also reasonably correlated with aboveground biomass estimated with tree measurements and allometric equations (see D, black line shows the LOESS regression between the two measures).

**Figure S5.** Spatial autocorrelation in the logged sectors of the canopy height models. Eighteen sectors (250 $km^2$) with large proportions of logged areas were first selection (A; bioclimatic domains are part of the Quebec's government ecological classification). For each sector separately, semi-variograms (B) were then produced with 10000 randomly selected pairs of 20 m × 20 m pixels (for each distance class) in the logged sectors of the canopy height models (P95 in meters). The red vertical bar in B) shows the 250 m minimum distance retained to select pixels of the training and validation datasets.

**Figure S6.** Height thresholds per bioclimatic domain that were retained to filter aberrant heights within the training and validation datasets. Bioclimatic domains (A) are part of the Quebec's government ecological classification. For each bioclimatic domain, the maximum height threshold per age class and per bioclimatic domain (curves in B) was then defined as the mean of individual species threshold (see Figure S4) weighted by their relative abundance in each bioclimatic domain. The curves in (B) correspond to the colors of bioclimatic domains in (A). Pixel color gradient in (B) shows the density of plots in the original dataset before filtering with height threshold, 10 450 plots out of a total of 248 969 were excluded from the original dataset.

**Figure S7.** Percentage of broadleaved trees used to document forest composition. The Laurentian Forestry Centre (Canadian Forest Service) established the proportion of deciduous trees with a Random Forest classification of the territory based on Landsat images between the provinces of Ontario and Newfoundland and Labrador (30 m × 30 m raster), and more than 10 k inventory plots from Quebec's government. The boreal zone for which forest growth was projected is delimited in by a red line.

**Figure S8.** Comparisons of model predictive performance with different combinations of temperature ($T_{MEAN}$, $T_{MAX}$) and climate moisture variables ($CMI_{GS}$, $CMI_{SM}$). For each combination, model predictive performance was evaluated as in Figure 2; by comparing observed and predicted heights (P95) in the validation dataset (20% of the entire dataset). Interactive effects of time-since-last clearcut, temperature, and climate moisture variables upon forest height were then evaluated as in Figure 3 (a random sampling of 999 observations in the original dataset was used to predict forest height across all one-year time-since-last clearcut classes). The fine grey lines show the entire variability of predicted heights without fixing any parameters. Colored lines and ribbons show the central tendency and 70% confidence intervals of predicted heights after fixing different parameters. Climate variables were fixed within their observed range in the original dataset and fixed values are shown in the upper left corners.

**Figure S9.** Correlation matrix between continuous variables within the training and validation datasets; Pearson's r is specified for each pair of variables.

**Figure S10.** Interactive effects of time-since-last clearcut, forest composition, and all other explanatory variables upon forest height. A unique random sampling of 999 observations in the original dataset was used to predict forest height across all one-year time-since-last clearcut classes. The fine grey lines show the whole variability of predicted heights without fixing any parameters. Colored lines and ribbons show the central tendency and 70% confidence intervals of predicted heights after fixing different parameters. Forest composition was fixed as coniferous, mixed, or broadleaved (i.e., 5%, 50%, and 95% of broadleaved trees, respectively). Other explanatory variables were fixed within their observed range in the original dataset (i.e., equal intervals between minimum and maximum observed values) and fixed values are shown in the upper left corners.

**Figure S11.** Predicted potential post-logging regrowth rates within the boreal zone. For each forest composition class (i.e., A: coniferous, B: mixed, or C: broadleaved), height growth rates in the first 50 years following clearcut was predicted following different combinations of climate scenarios (see Figure S1). The different temperature ($T_{MEAN}$) scenarios are displayed along the horizontal line (Tbse: Baseline 1980-2020, T4.5: RCP4.5 2041-2070 and T8.5: RCP8.5 2041-2070). Similarly, the different climate moisture scenarios ($CMI_{SM}$) are displayed along the horizontal line (Mbse: Baseline 1980-2020, M4.5: RCP4.5 2041-2070 and M8.5: RCP8.5 2041-2070). Baselines growth rates (i.e., $T_{MEAN}$ and $CMI_{SM}$ for the 1980-2020 period) are thus shown in the first column and row of each composition class (i.e., A, B and C). Changes in height growth rates with different combinations of projected climate scenarios (2041-2070) are displayed as percentage of changes from baseline scenarios. Finally, for each composition class (A, B, and C), model prediction uncertainties are shown for the three different climate scenarios (i.e., Baseline, RCP4.5, and RCP8.5).

**Figure S12.** Enlarged versions of the histograms in Figure S11 (Predicted potential post-logging regrowth rates within the boreal zone). For each composition class (A, B, and C), the different temperature ($T_{MEAN}$) scenarios are displayed along the horizontal line (Tbse: Baseline 1980-2020, T4.5: RCP4.5 2041-2070 and T8.5: RCP8.5 2041-2070). Similarly, the different climate moisture scenarios ($CMI_{SM}$) are displayed along the horizontal line (Mbse: Baseline 1980-2020, M4.5: RCP4.5 2041-2070 and M8.5: RCP8.5 2041-2070). Baselines growth rates (i.e., $T_{MEAN}$ and $CMI_{SM}$ for the 1980-2020 period) are thus shown in the first column and row of each composition class (i.e., A, B and C). Changes in height growth rates with different combinations of projected climate scenarios (2041-2070) are displayed as percentage of changes from baseline scenarios.

**Figure S13.** Forest growth rates for the different combinations of climate scenarios. For each forest composition class, height growth rates in the first 50 years following clearcut were predicted following different combinations of climate scenarios (see Figure S1). In scenario names, *T* and *M* respectively indicate temperature and moisture, while *bse*, *4.5*, and *8.5* indicate baselines, RCP 4.5 and 8.5 scenarios, respectively. Forest composition classes are displayed as colored lines (i.e., C: coniferous, M: mixed, and B: broadleaved). Density curves were obtained with a random sampling of 5% of 20 m pixels within each raster showed in Figure S11.

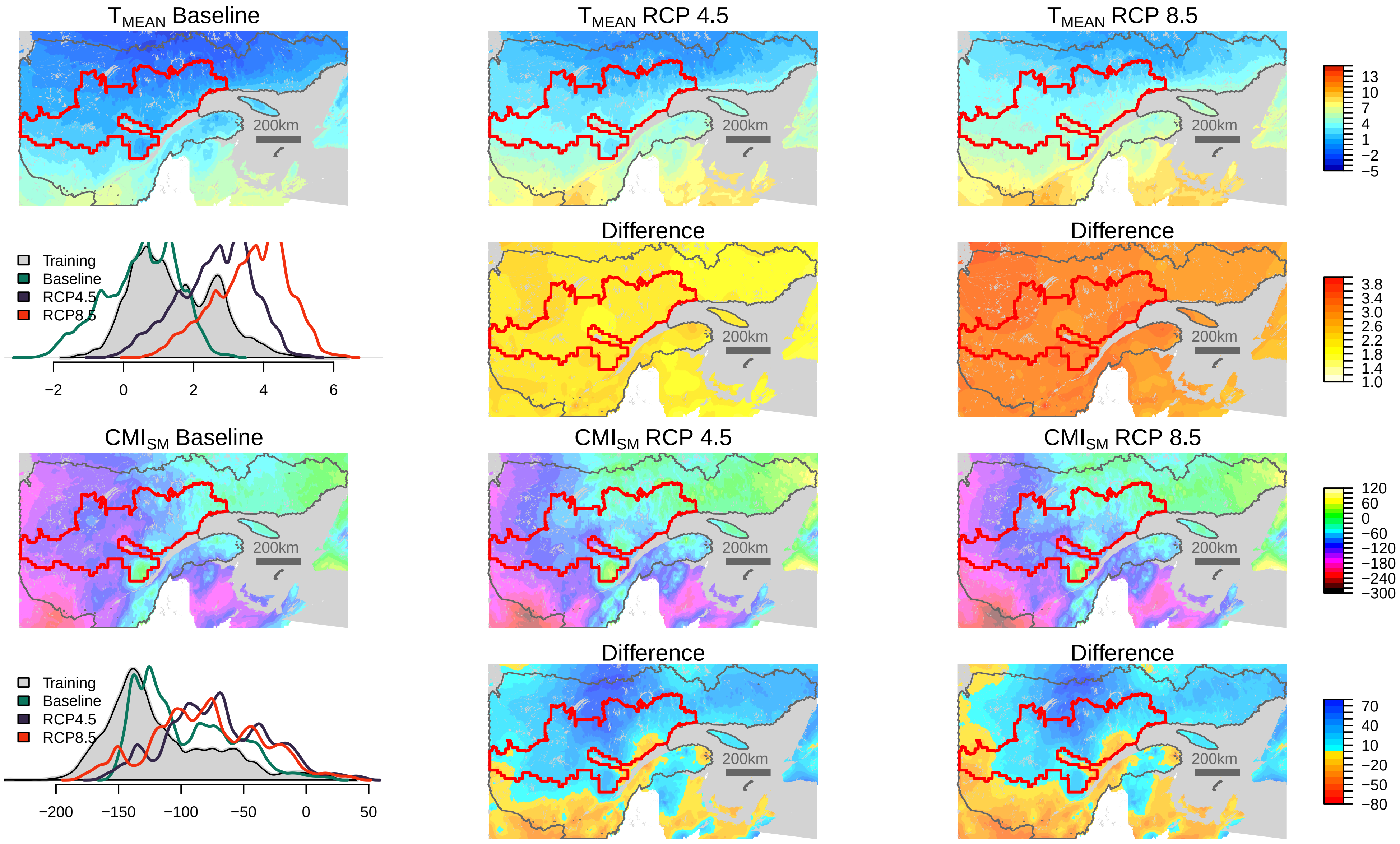

$T_{MEAN}$ Baseline
$T_{MEAN}$ RCP 4.5
$T_{MEAN}$ RCP 8.5
200km
13
10
7
4
1
−2
−5
Training
Baseline
RCP4.5
RCP8.5
−2
0
2
4
6
Difference
Difference
3.8
3.4
3.0
2.6
2.2
1.8
1.4
1.0
$CMI_{SM}$ Baseline
$CMI_{SM}$ RCP 4.5
$CMI_{SM}$ RCP 8.5
120
60
0
−60
−120
−180
−240
−300
−200
−150
−100
−50
0
50
70
40
10
−20
−50
−80


Figure S1

A. Elevation (meters)

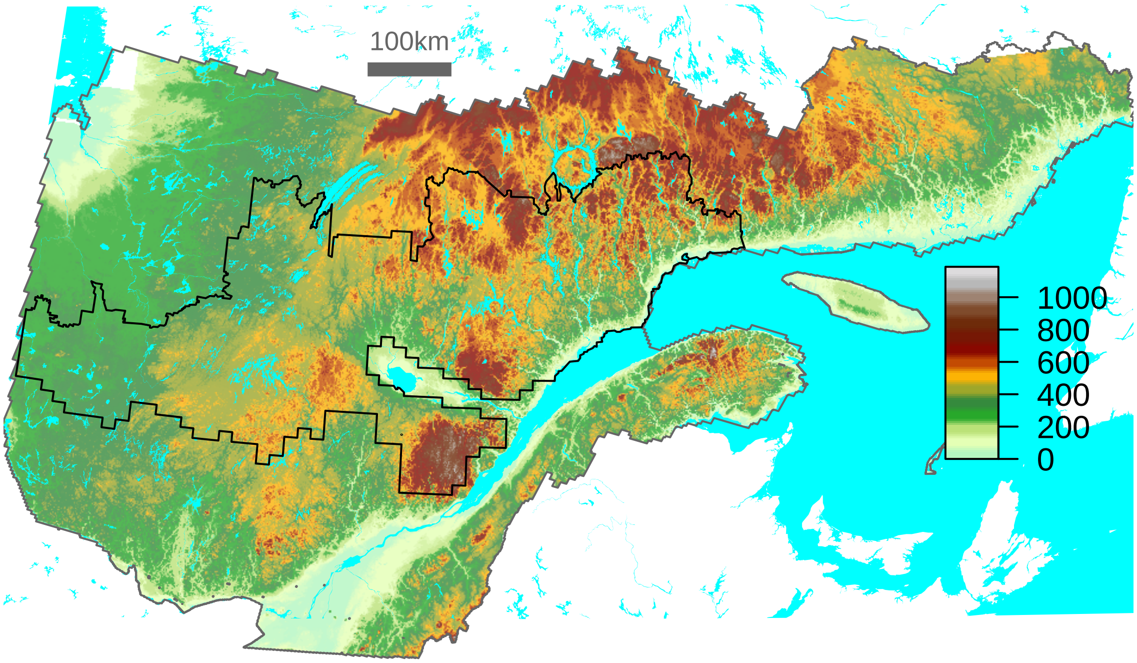


B. Surface deposits categories

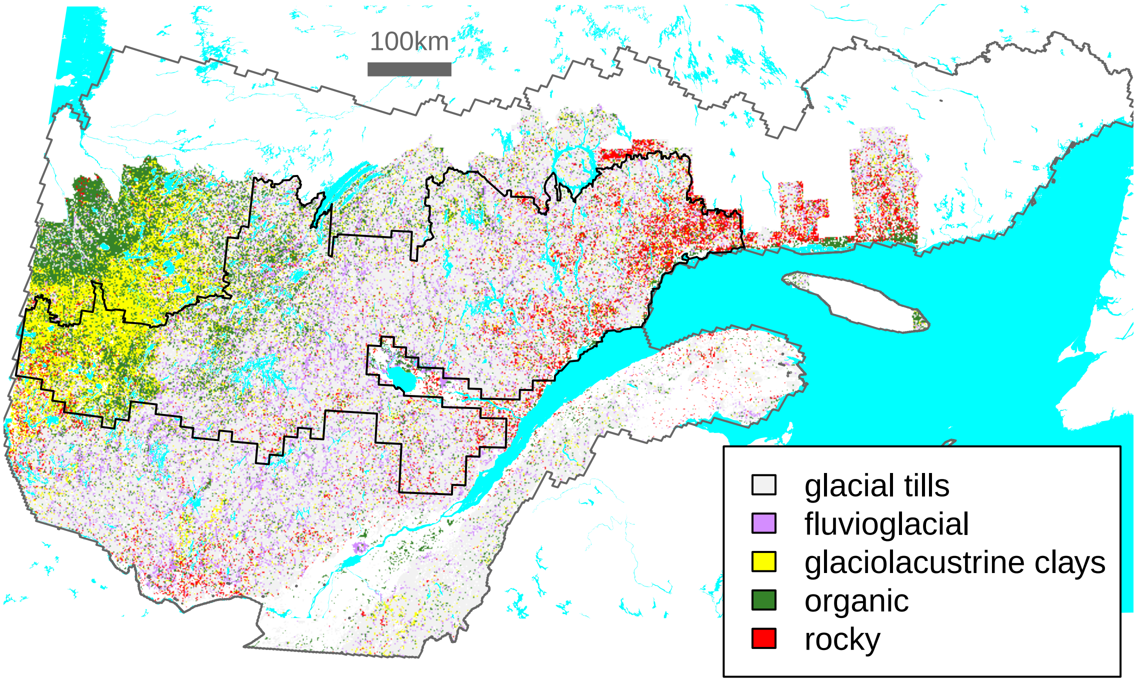

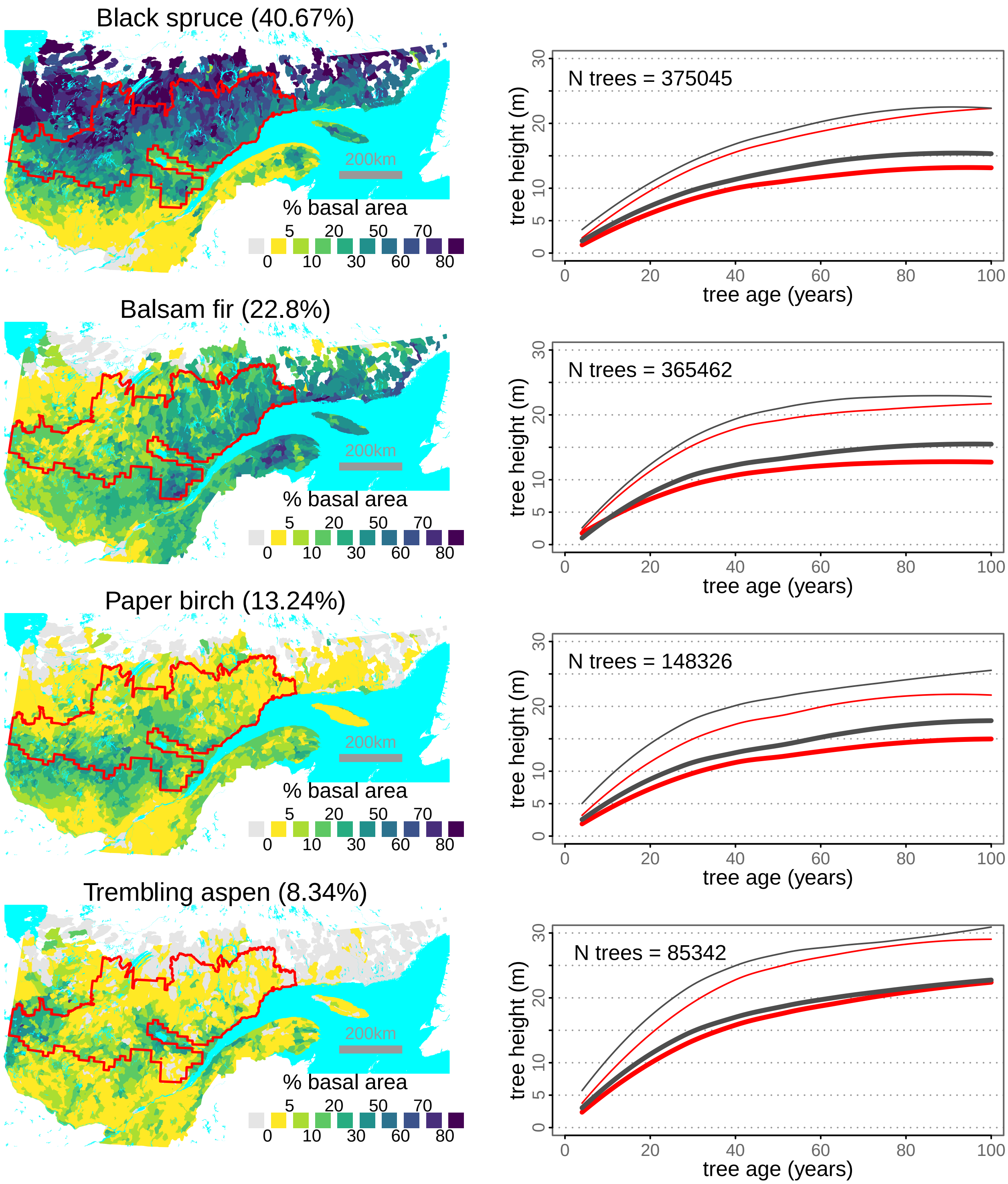
Black spruce (40.67%)
200km
% basal area
5 20 50 70
0 10 30 60 80
N trees = 375045
tree height (m)
tree age (years)
Balsam fir (22.8%)
200km
% basal area
5 20 50 70
0 10 30 60 80
N trees = 365462
tree height (m)
tree age (years)
Paper birch (13.24%)
200km
% basal area
5 20 50 70
0 10 30 60 80
N trees = 148326
tree height (m)
tree age (years)
Trembling aspen (8.34%)
200km
% basal area
5 20 50 70
0 10 30 60 80
N trees = 85342
tree height (m)
tree age (years)

Figure S3

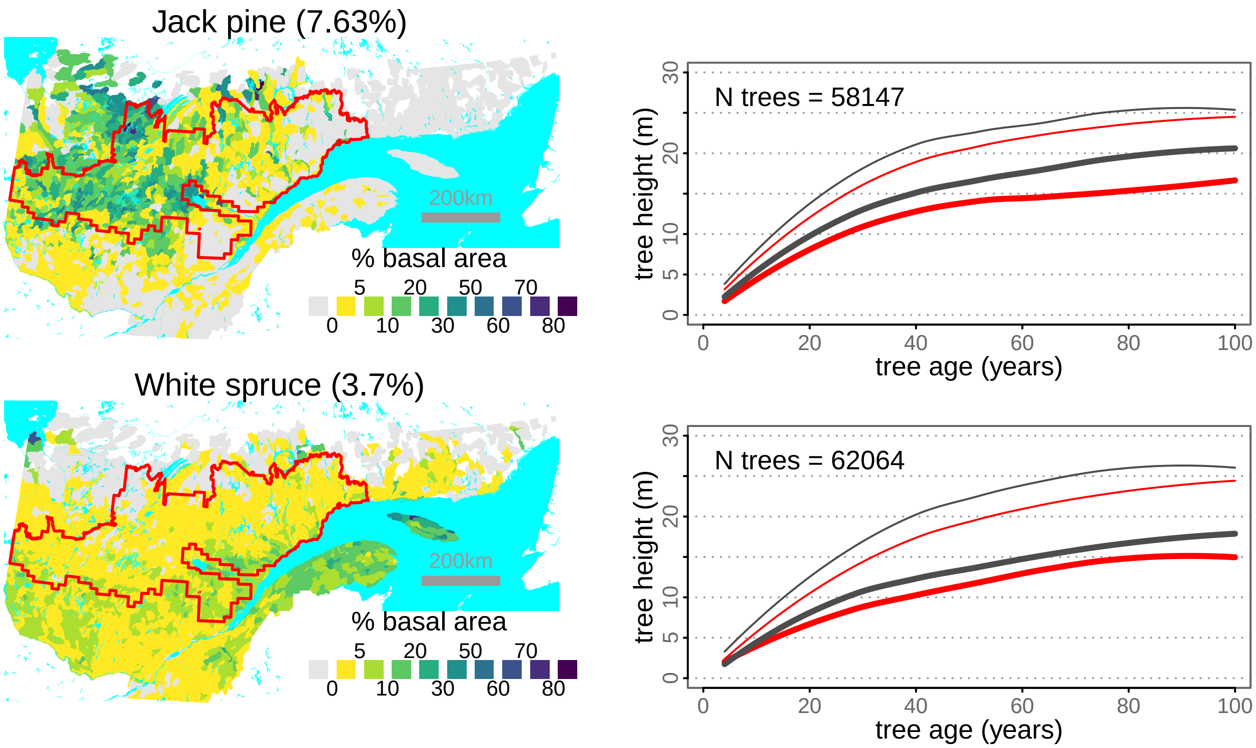
Jack pine (7.63%)
200km
% basal area
5 20 50 70
0 10 30 60 80
N trees = 58147
tree height (m)
tree age (years)
White spruce (3.7%)
200km
% basal area
5 20 50 70
0 10 30 60 80
N trees = 62064
tree height (m)
tree age (years)

Figure S3

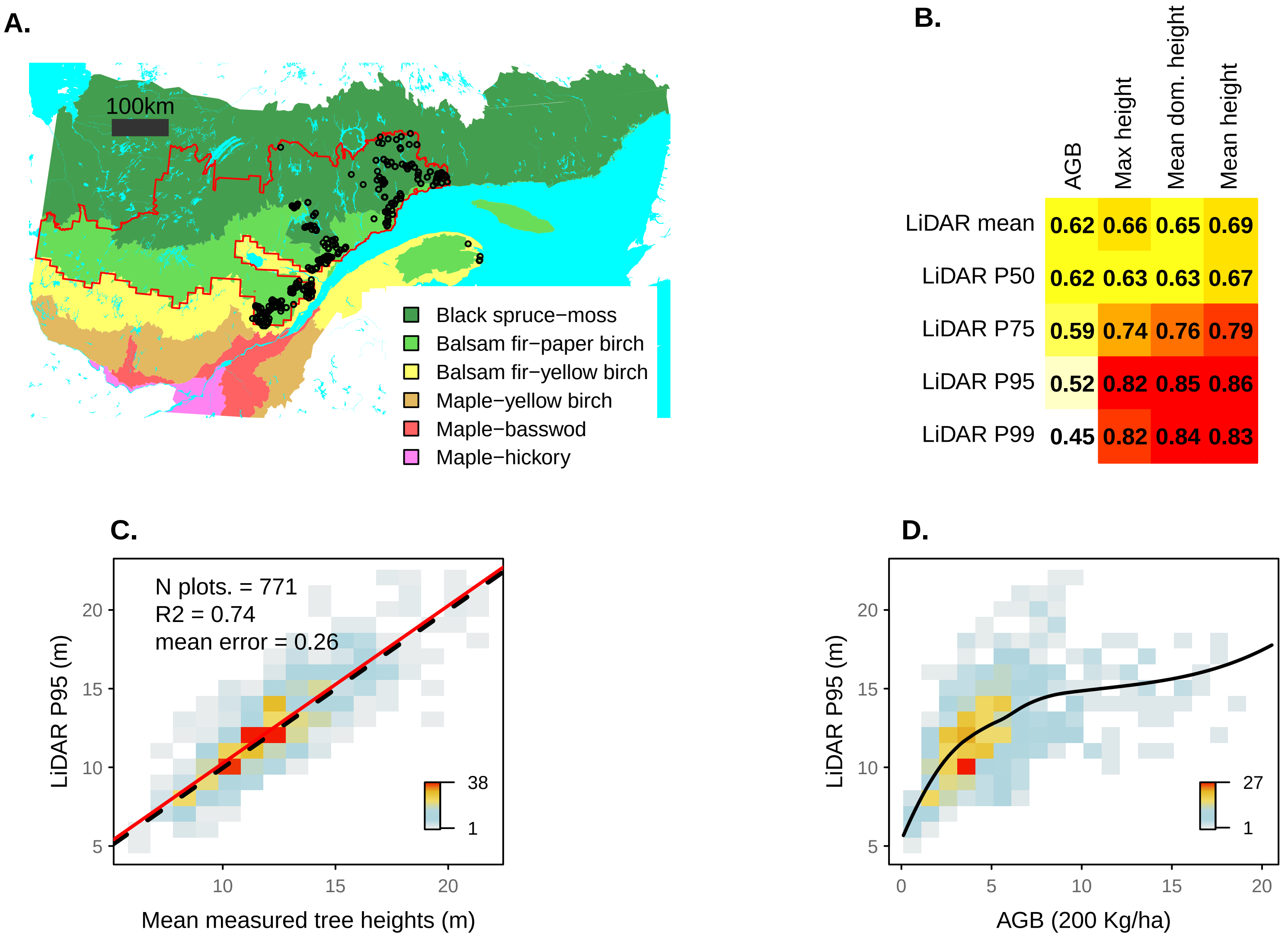
A.
100km
Black spruce−moss
Balsam fir−paper birch
Balsam fir−yellow birch
Maple−yellow birch
Maple−basswod
Maple−hickory
B.
AGB
Max height
Mean dom. height
Mean height
LiDAR mean 0.62 0.66 0.65 0.69
LiDAR P50 0.62 0.63 0.63 0.67
LiDAR P75 0.59 0.74 0.76 0.79
LiDAR P95 0.52 0.82 0.85 0.86
LiDAR P99 0.45 0.82 0.84 0.83
C.
N plots. = 771
R2 = 0.74
mean error = 0.26
38
1
LiDAR P95 (m)
Mean measured tree heights (m)
D.
27
1
LiDAR P95 (m)
AGB (200 Kg/ha)

Figure S4

**A. Sampled sectors**

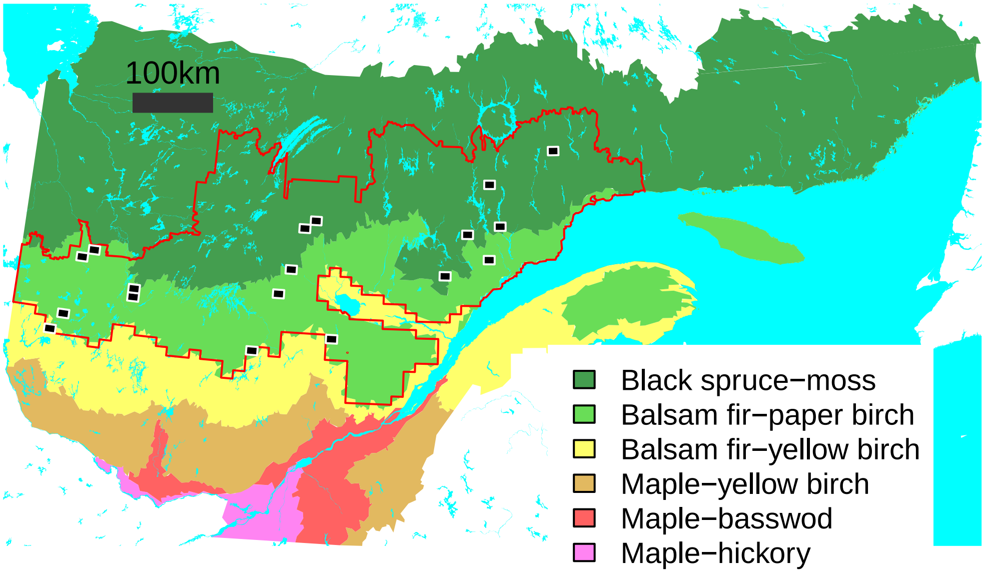


**B. Semivariograms**

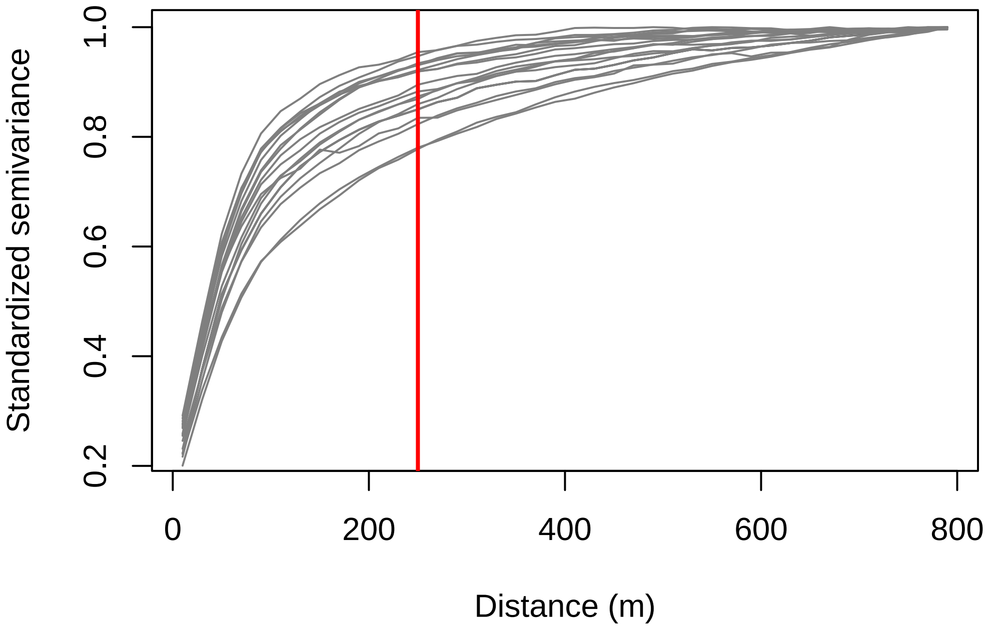

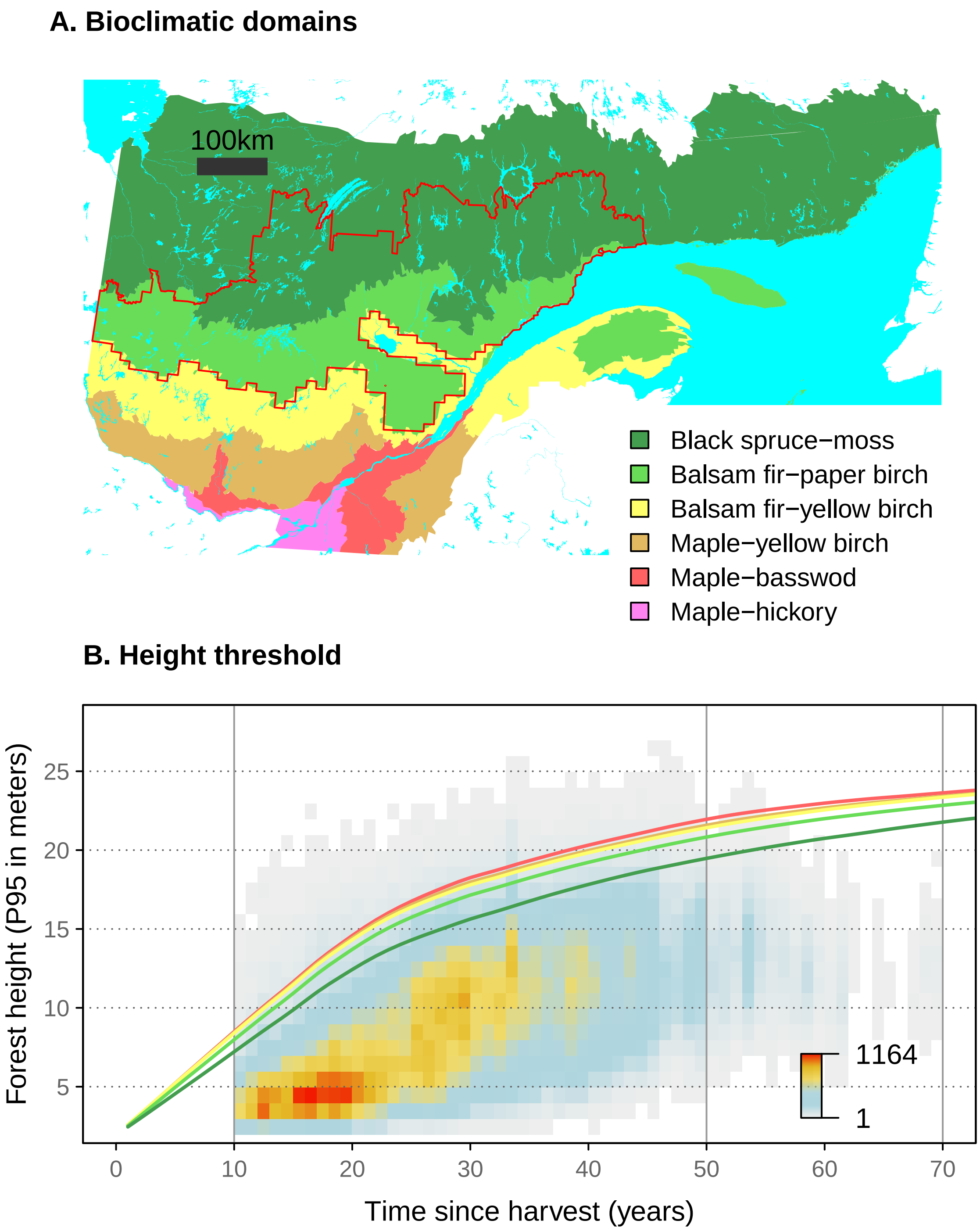

A. Bioclimatic domains
100km
Black spruce−moss
Balsam fir−paper birch
Balsam fir−yellow birch
Maple−yellow birch
Maple−basswod
Maple−hickory
B. Height threshold
Forest height (P95 in meters)
25
20
15
10
5
1164
1
0
10
20
30
40
50
60
70
Time since harvest (years)


Figure S6

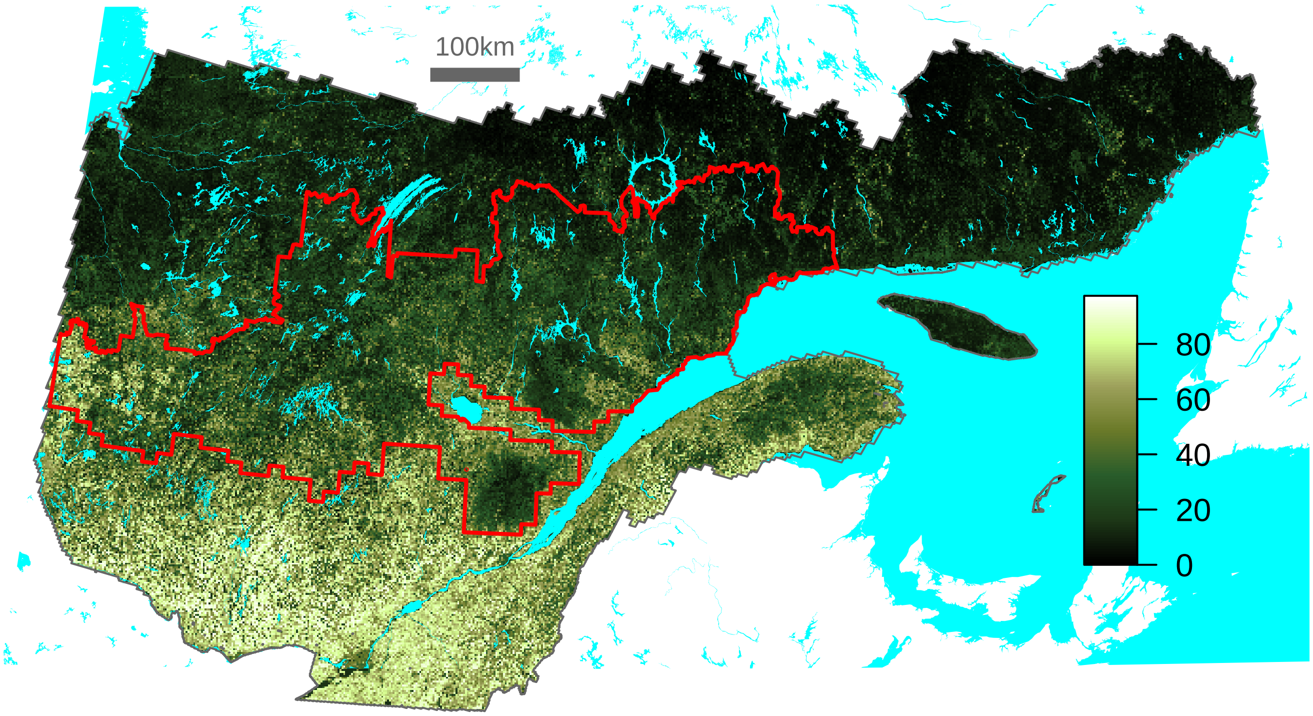


Figure S7

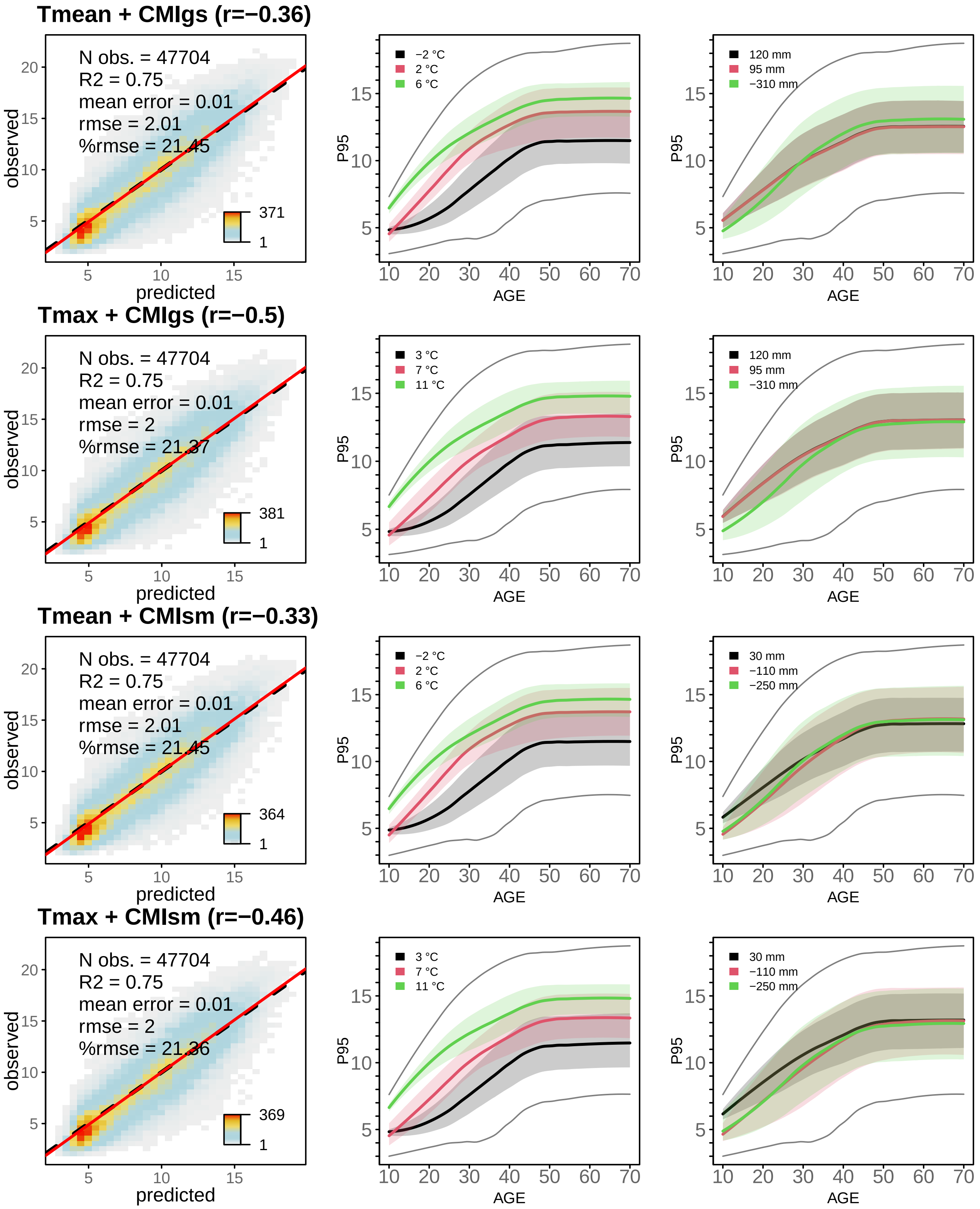
Tmean + CMIgs (r=−0.36)
N obs. = 47704
R2 = 0.75
mean error = 0.01
rmse = 2.01
%rmse = 21.45
371
1
observed
predicted
−2 °C
2 °C
6 °C
120 mm
95 mm
−310 mm
P95
AGE
Tmax + CMIgs (r=−0.5)
N obs. = 47704
R2 = 0.75
mean error = 0.01
rmse = 2
%rmse = 21.37
381
1
3 °C
7 °C
11 °C
120 mm
95 mm
−310 mm
Tmean + CMIsm (r=−0.33)
N obs. = 47704
R2 = 0.75
mean error = 0.01
rmse = 2.01
%rmse = 21.45
364
1
−2 °C
2 °C
6 °C
30 mm
−110 mm
−250 mm
Tmax + CMIsm (r=−0.46)
N obs. = 47704
R2 = 0.75
mean error = 0.01
rmse = 2
%rmse = 21.36
369
1
3 °C
7 °C
11 °C
30 mm
−110 mm
−250 mm

Figure S8

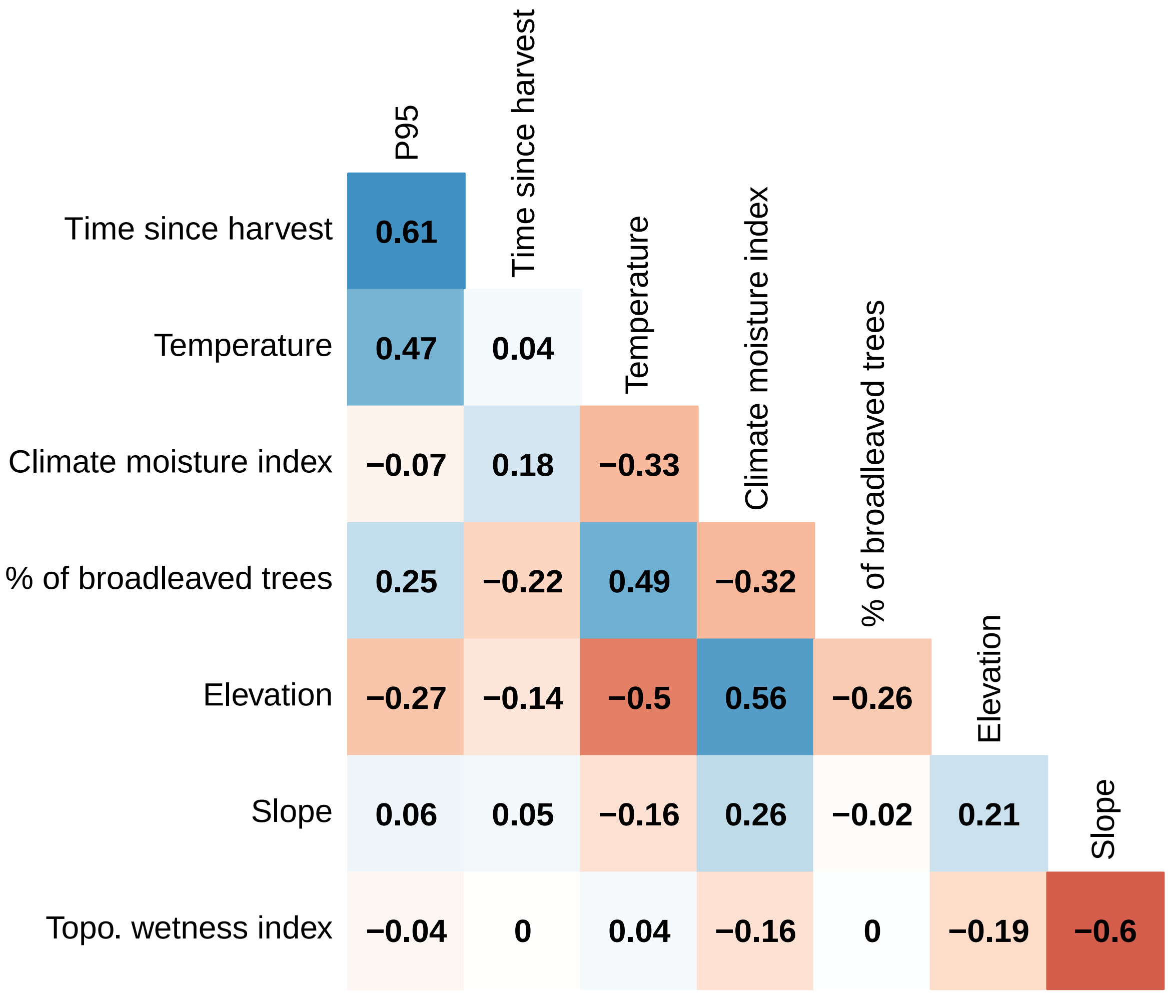
P95
Time since harvest
Temperature
Climate moisture index
% of broadleaved trees
Elevation
Slope
Time since harvest 0.61
Temperature 0.47 0.04
Climate moisture index −0.07 0.18 −0.33
% of broadleaved trees 0.25 −0.22 0.49 −0.32
Elevation −0.27 −0.14 −0.5 0.56 −0.26
Slope 0.06 0.05 −0.16 0.26 −0.02 0.21
Topo. wetness index −0.04 0 0.04 −0.16 0 −0.19 −0.6

Figure S9

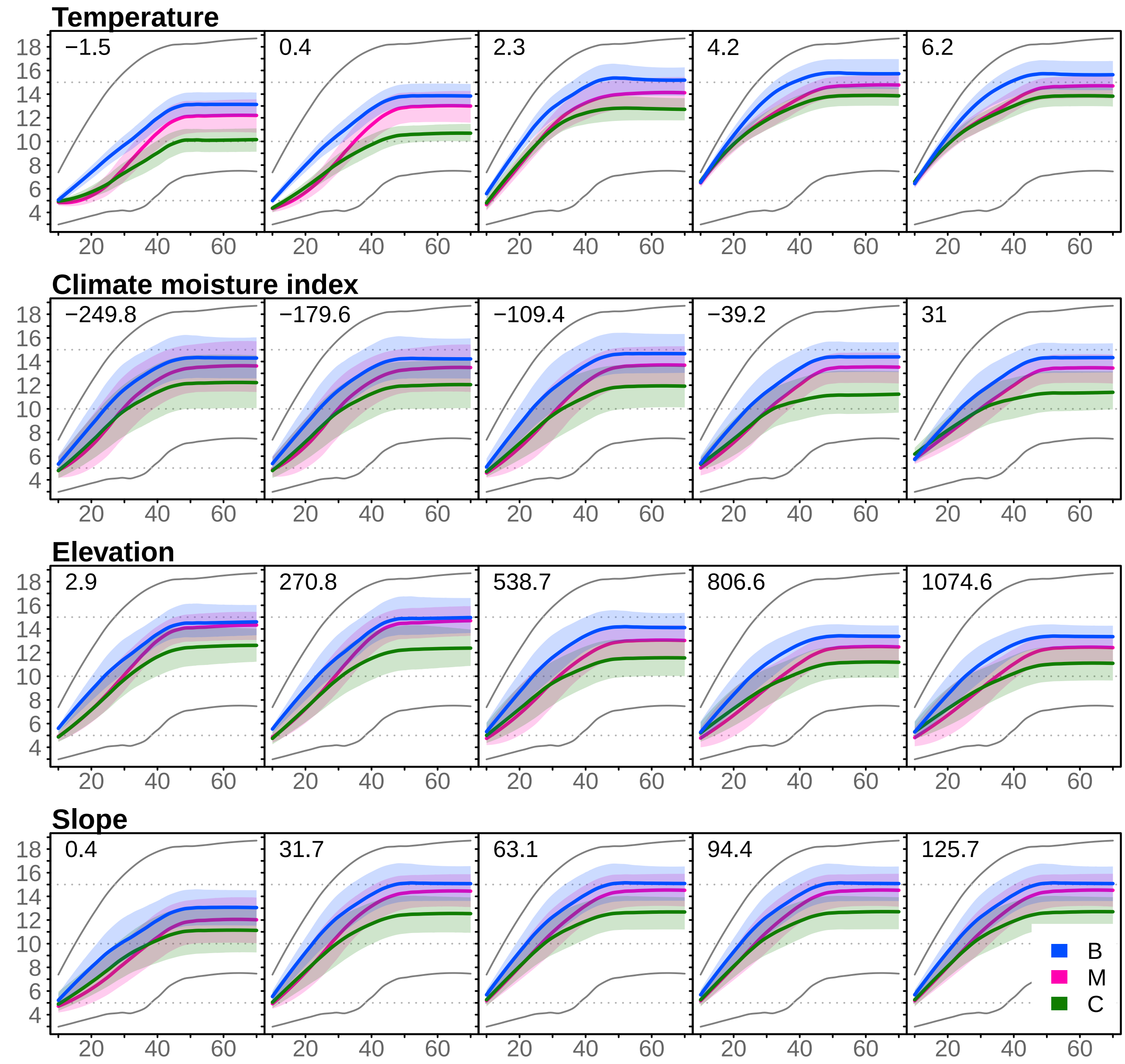
Temperature
−1.5
0.4
2.3
4.2
6.2
Climate moisture index
−249.8
−179.6
−109.4
−39.2
31
Elevation
2.9
270.8
538.7
806.6
1074.6
Slope
0.4
31.7
63.1
94.4
125.7
18
16
14
12
10
8
6
4
20
40
60
B
M
C

Figure S10

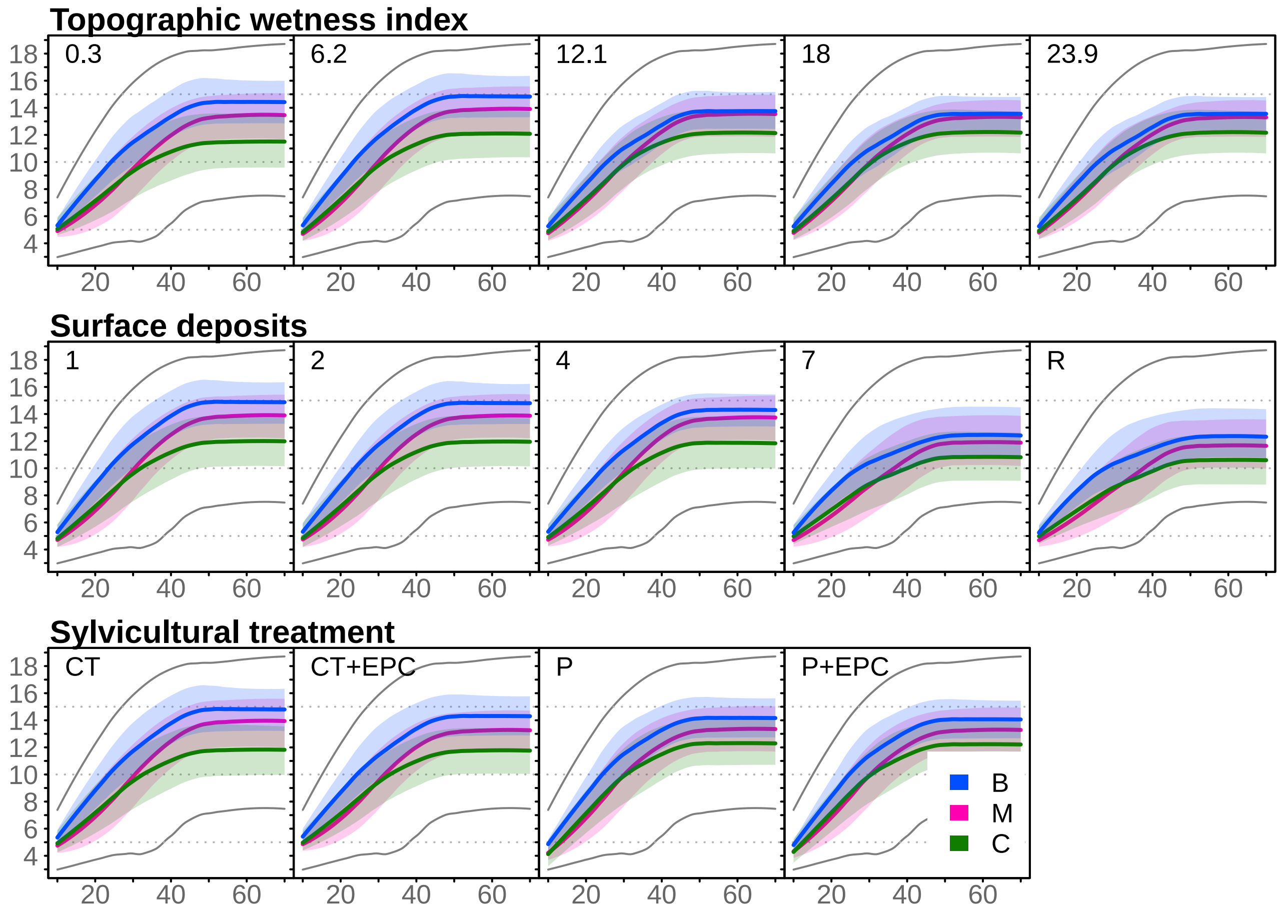

Topographic wetness index
0.3
6.2
12.1
18
23.9
Surface deposits
1
2
4
7
R
Sylvicultural treatment
CT
CT+EPC
P
P+EPC
B
M
C


Figure S10

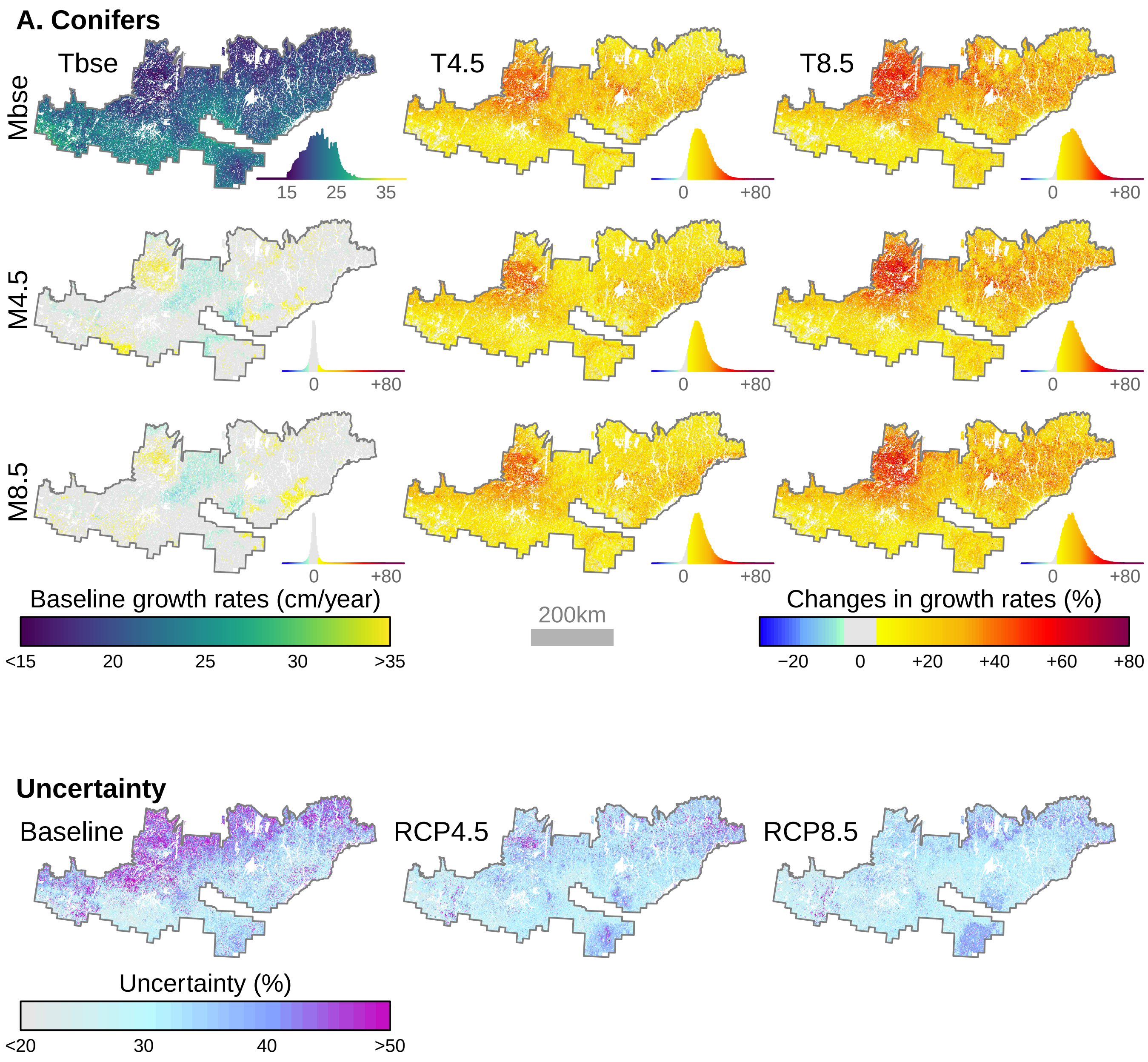
A. Conifers
Tbse
T4.5
T8.5
Mbse
M4.5
M8.5
15
25
35
0
+80
Baseline growth rates (cm/year)
<15
20
25
30
>35
200km
Changes in growth rates (%)
−20
0
+20
+40
+60
+80
Uncertainty
Baseline
RCP4.5
RCP8.5
Uncertainty (%)
<20
30
40
>50

Figure S11

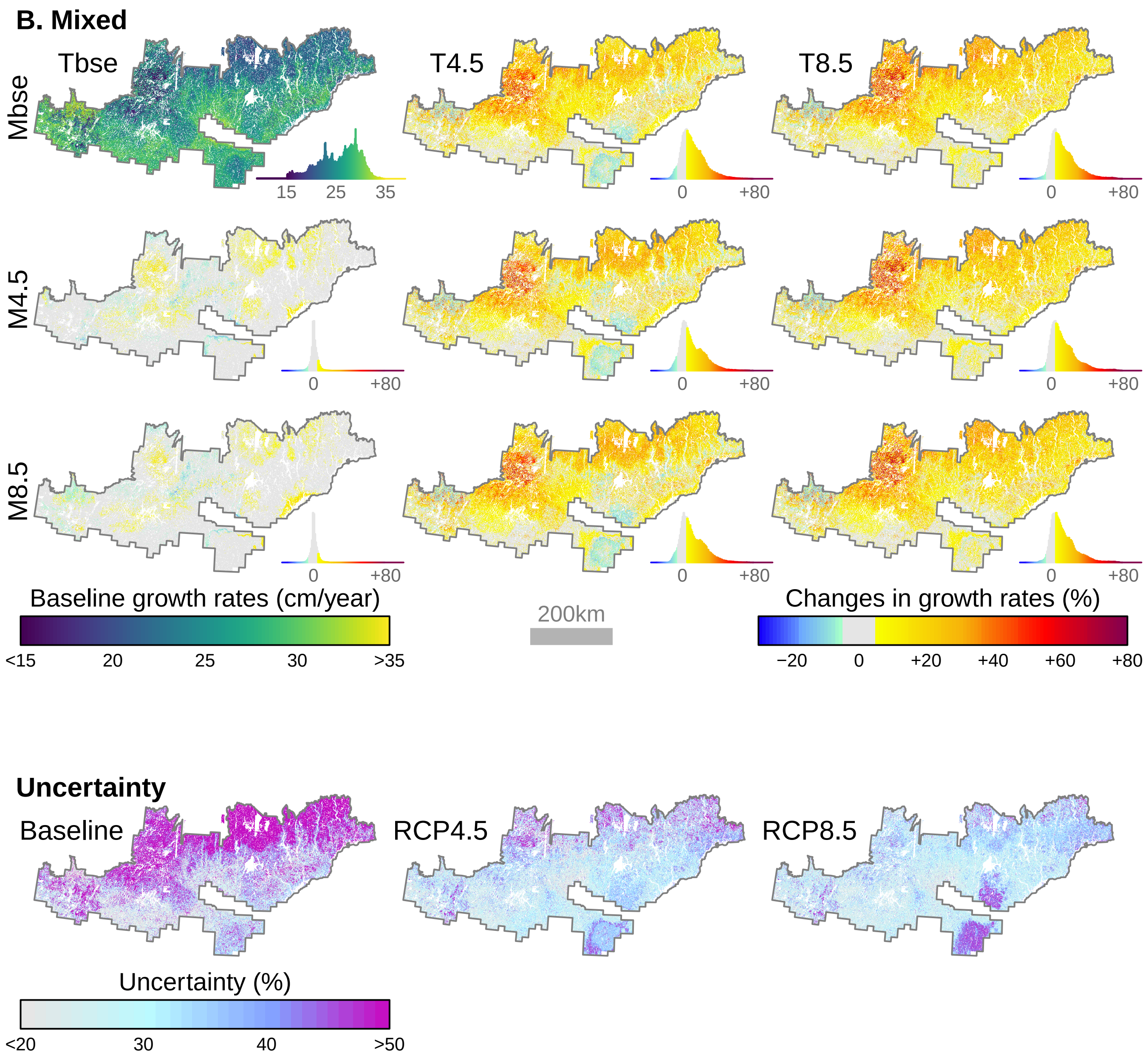
B. Mixed
Tbse
T4.5
T8.5
Mbse
M4.5
M8.5
15
25
35
0
+80
Baseline growth rates (cm/year)
<15
20
25
30
>35
200km
Changes in growth rates (%)
−20
0
+20
+40
+60
+80
Uncertainty
Baseline
RCP4.5
RCP8.5
Uncertainty (%)
<20
30
40
>50

Figure S11

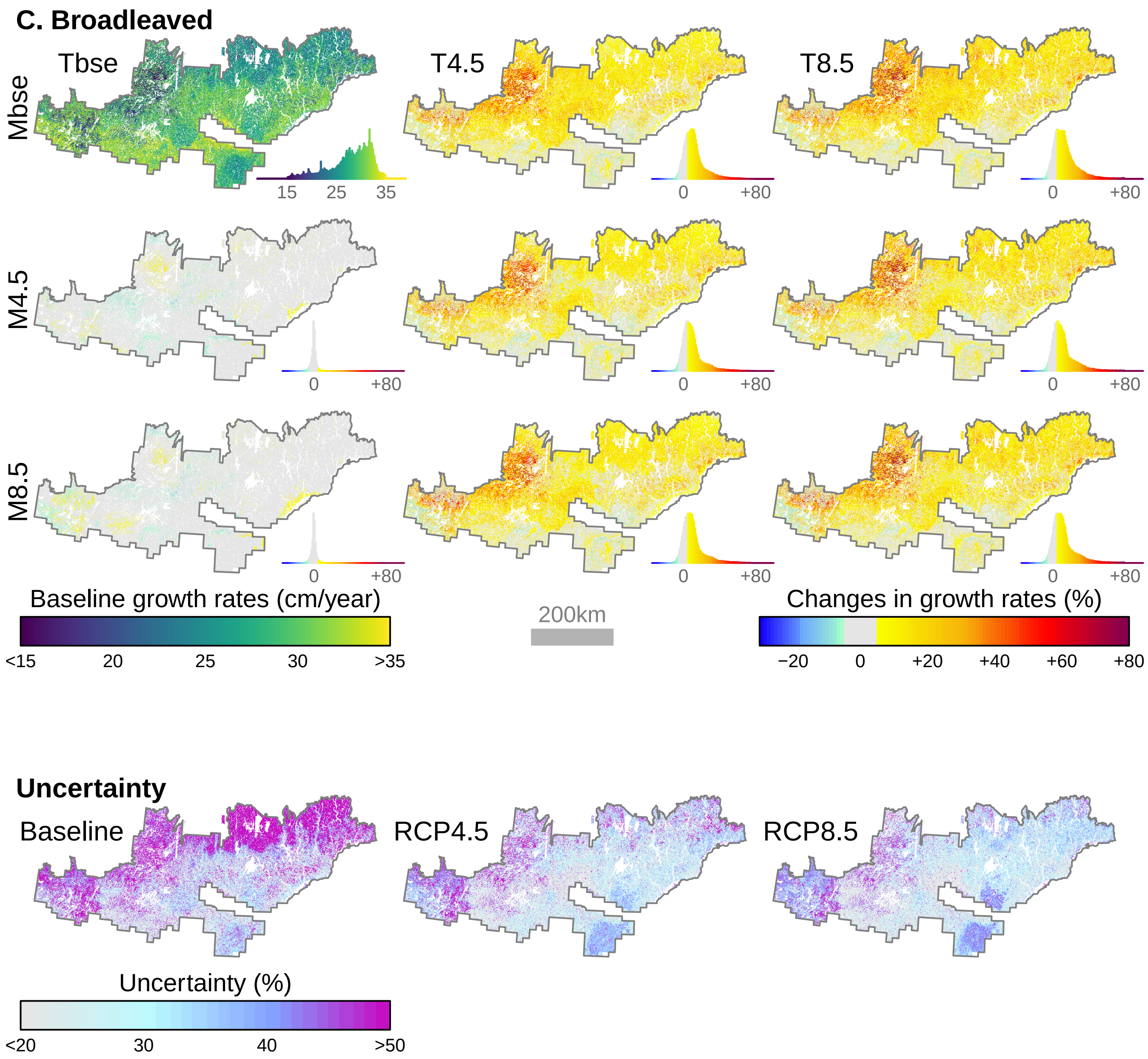

C. Broadleaved
Tbse
T4.5
T8.5
Mbse
M4.5
M8.5
15
25
35
0
+80
Baseline growth rates (cm/year)
<15
20
25
30
>35
200km
Changes in growth rates (%)
−20
0
+20
+40
+60
+80
Uncertainty
Baseline
RCP4.5
RCP8.5
Uncertainty (%)
<20
30
40
>50


Figure S11

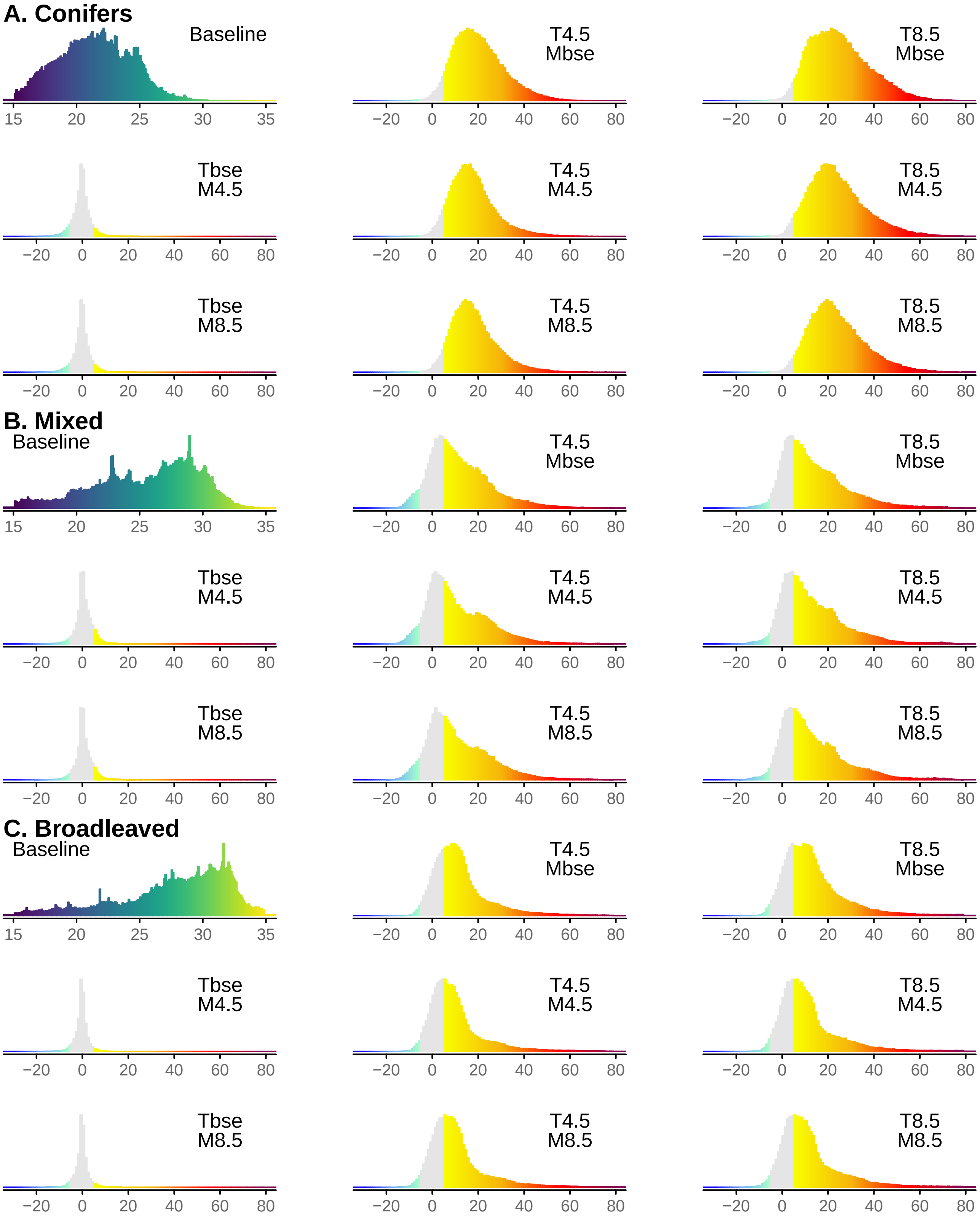
A. Conifers
Baseline
T4.5 Mbse
T8.5 Mbse
Tbse M4.5
T4.5 M4.5
T8.5 M4.5
Tbse M8.5
T4.5 M8.5
T8.5 M8.5
B. Mixed
Baseline
C. Broadleaved
15 20 25 30 35
−20 0 20 40 60 80

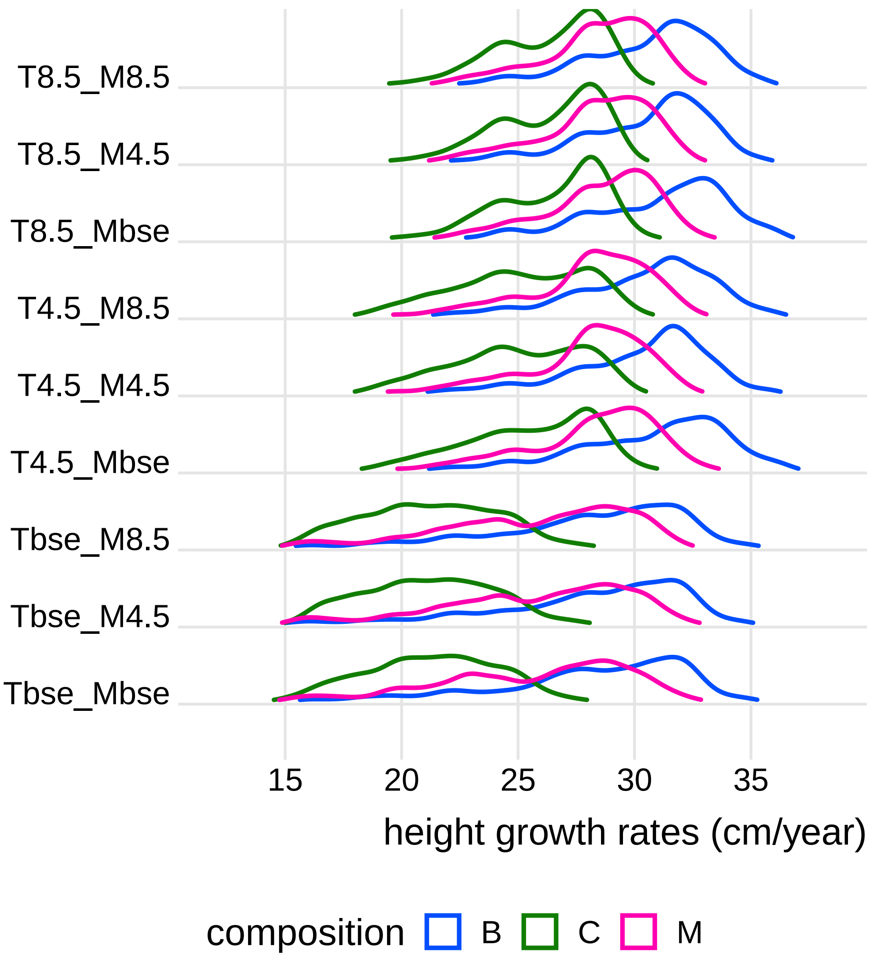
T8.5_M8.5
T8.5_M4.5
T8.5_Mbse
T4.5_M8.5
T4.5_M4.5
T4.5_Mbse
Tbse_M8.5
Tbse_M4.5
Tbse_Mbse
15
20
25
30
35
height growth rates (cm/year)
composition
B
C
M
